# Data-driven dynamical modelling of a pathogen-infected plant gene regulatory network: a comparative analysis

**DOI:** 10.1101/2022.02.03.479002

**Authors:** Mathias Foo, Leander Dony, Fei He

## Abstract

Recent advances in synthetic biology have enabled the design of genetic feedback control circuits that could be implemented to build resilient plants against pathogen attacks. To facilitate the proper design of these genetic feedback control circuits, an accurate model that is able to capture the vital dynamical behaviour of the pathogen-infected plant is required. In this study, using a data-driven modelling approach, we develop and compare four dynamical models (i.e. linear, Michaelis-Menten, standard S-System and extended S-System) of a pathogen-infected plant gene regulatory network (GRN). These models are then assessed across several criteria, i.e. ease of identifying the type of gene regulation, the predictive capability, Akaike Information Criterion (AIC) and the robustness to parameter uncertainty to determine its viability of modelling the pathogen-infected plant GRN. Using our defined ranking score, our analyses show that while the extended S-System model ranks highest in the overall comparison, the performance of the linear model is more consistent throughout the comparison, making it the preferred model for this pathogen-infected plant GRN.

## I. INTRODUCTION

One of the common fungal pathogens that infects plant is the *Botrytis cinerea*. When infection occurs, the interactions between the pathogen and the host plant often lead to the host plant succumb to diseases. This is because pathogen often disrupts the host defense mechanism through secretion of a range of proteins, small RNAs and metabolites to aid their colonisation [1–4]. Advances in the area of molecular biology have provided plant synthetic biologists means of improving plant resilience through the use of synthetic feedback control circuits (see *e.g.* [5]) to restore the regulation that is affected by the pathogen attack [6]. Pathogen affected genes involved in defence tend to have their expression levels compromised, leading to their reduced functional ability [7, 8]. The synthetic feedback control circuits would sense the changes in the expression level of pathogen affected genes, where the genes cis-regulatory elements are modified resulting in changes in their regulations and expression levels (see [9] and references therein) and regulate appropriate transcription factor to allow the compromised expression levels to be controlled thereby enabling plant to recover their defence functionality.

To facilitate the design of these synthetic feedback control circuits, an accurate dynamical model depicting the gene regulatory network (GRN) involved in the plant defense mechanism is required. In our previous study [6], equipped with the temporal data of gene expressions [10] and the knowledge of the interacting genes involved in plant defence [9], a linear dynamical model is developed using a data-driven modelling approach to model the pathogen-infected plant GRN with good accuracy and subsequently used to design and develop a framework of engineering resilience plant using synthetic genetic feedback control circuits. In this study, as a follow up to [6], we aim to answer the following question: *“When using the data-driven modelling approach, given the temporal data and knowledge about the pathogen-infected plant GRN interaction, is the linear model the most viable model to facilitate the design of synthetic feedback control and if not what is the alternate candidate model*?”

Since the advancement in the area of Systems Biology, GRN modelling has been extensively studied (see the review paper by [11] and references therein). According to [11], most of the models described in those studies can be categorised into four main classes in the order of increased complexity — part list model (e.g. description of the GRN component), topology model (e.g. directed graph model), control logic model (*e.g.* Boolean function model) and dynamical model (e.g. differential equation model). The models developed here are often based on first principles, *i.e.* using the biological understanding of the interacting components.

With the access to high throughput data at molecular level becoming available, attention turns to another branch of modelling approach called reverse engineering [12, 13], where models are developed in the attempt to fit those data using various methods such as correlation-based method, Bayesian networks, regression analysis, information theoretical approaches, Gaussian graphical models, dynamic differential equations, etc [14–17]. In a reverse engineering approach, usually there is no assumption about the model structure and the interacting components. The development in this area often parallels the development of GRN network inference algorithms, where the types and directions of regulation between components are inferred directly from data (see the review paper [18] and references therein). As a note, in the area of systems and control engineering [19], the reverse engineering approach is also known as system identification or data-driven modelling.

Here, we would like to make several remarks to provide readers the main scope of this study. First, this study is not about comparing network inference algorithms, hence the discussion on this topic is beyond the scope of this study. Interested readers can see the following review papers [20, 21] for more details. Second, unlike typical reverse engineering (data-driven modellling) approaches that assume almost no prior knowledge about the GRNs and the model structures, here we have some knowledge about the interacting genes and we have a set of candidate model structures of interest to be compared. Thus, our ‘network inference’ approach will be simpler with the focus on identifying the regulation type. Third, our study is system specific, *i.e.* a pathogen-infected plant GRN, and the main goal is to answer the key question posted above, i.e. the suitability of the linear dynamical model [6] in modelling a pathogen-infected plant GRN.

To the best of our knowledge, the only comparative study of different dynamical models for plant-specific GRN has been carried out in [22], where the authors compared several dynamical models for the GRN involved in plant flowering time. Different from that study, our study focuses on the data-driven modelling approach and uses different quantitative metrics for model comparison. In our comparative analysis, in addition to the linear model given in [6], we consider the Michaelis-Menten model and two S-System based models. The choice of these three models are motivated by their capabilities in modelling GRN (see *e.g.* [23–25]).

The manuscript is organised in the following manner. In Section II, we present the pathogen-infected plant GRN used as our case study. The main results on the comparative analysis of the four GRN models are presented and discussed in details in Section III. Finally, the discussion and conclusions are provided in Section IV.

## II. SYSTEM DESCRIPTION

The plant GRN involved in the defence against pathogen attack and used in this study is adapted from [10], where a subnetwork of nine genes — hereinafter termed 9GRN [6] — has been identified to be involved in the defence against *Botrytis cinerea,* as shown in Fig. 1. In Fig. 1, while the direction of regulation between genes in 9GRN is known, the type of regulation (i.e. activation or inhibition) is not entirely known. Among these nine genes, seven of them are directly affected by the pathogen, as indicated by the yellow hexagon. *CHE* and *ATML1* are part of the circadian clock genes as their oscillatory profiles are influenced by external light, as indicated by the red lightning. Moreover, the gene *CHE* has been identified to be an important gene in the plant defence mechanism and when it is affected by the pathogen, its expression level would decrease thereby reducing its defence capability [9, 10]. Therefore, it is imperative that the expression level of *CHE* being kept high and thus the role of the synthetic feedback control circuitry is to ensure its expression level stay high when under pathogen attack.

**FIG. 1:**
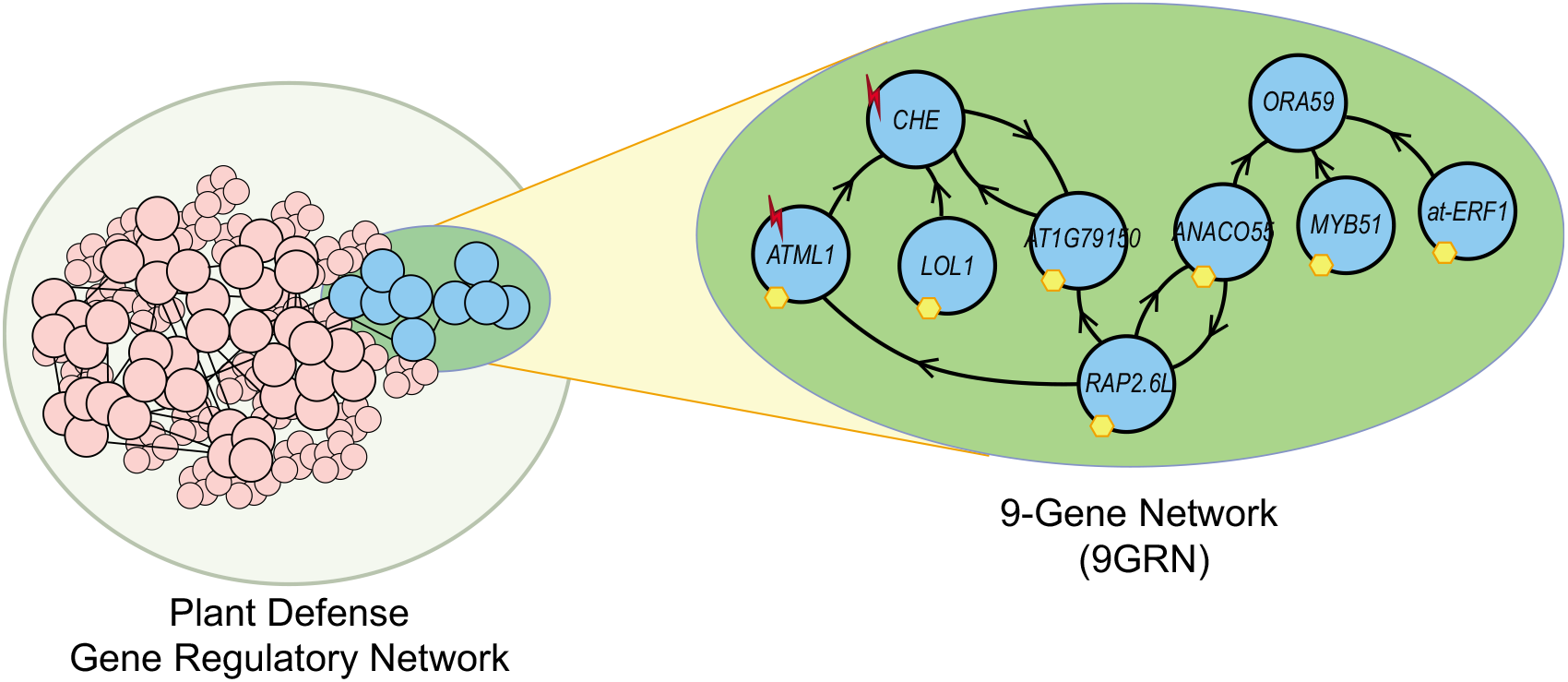
Plant (Arabidopsis) gene regulatory network (termed 9GRN) involved in the defence response to *Botrytis cinerea* adapted from [6]. The yellow hexagon symbol represents genes that have been identified to be directly affected by *Botrytis cinerea.* Red lightning symbol represents genes that are light regulated. The directional arrows indicate the influence of one gene to another despite its regulation type unknown.

**FIG. 2:**
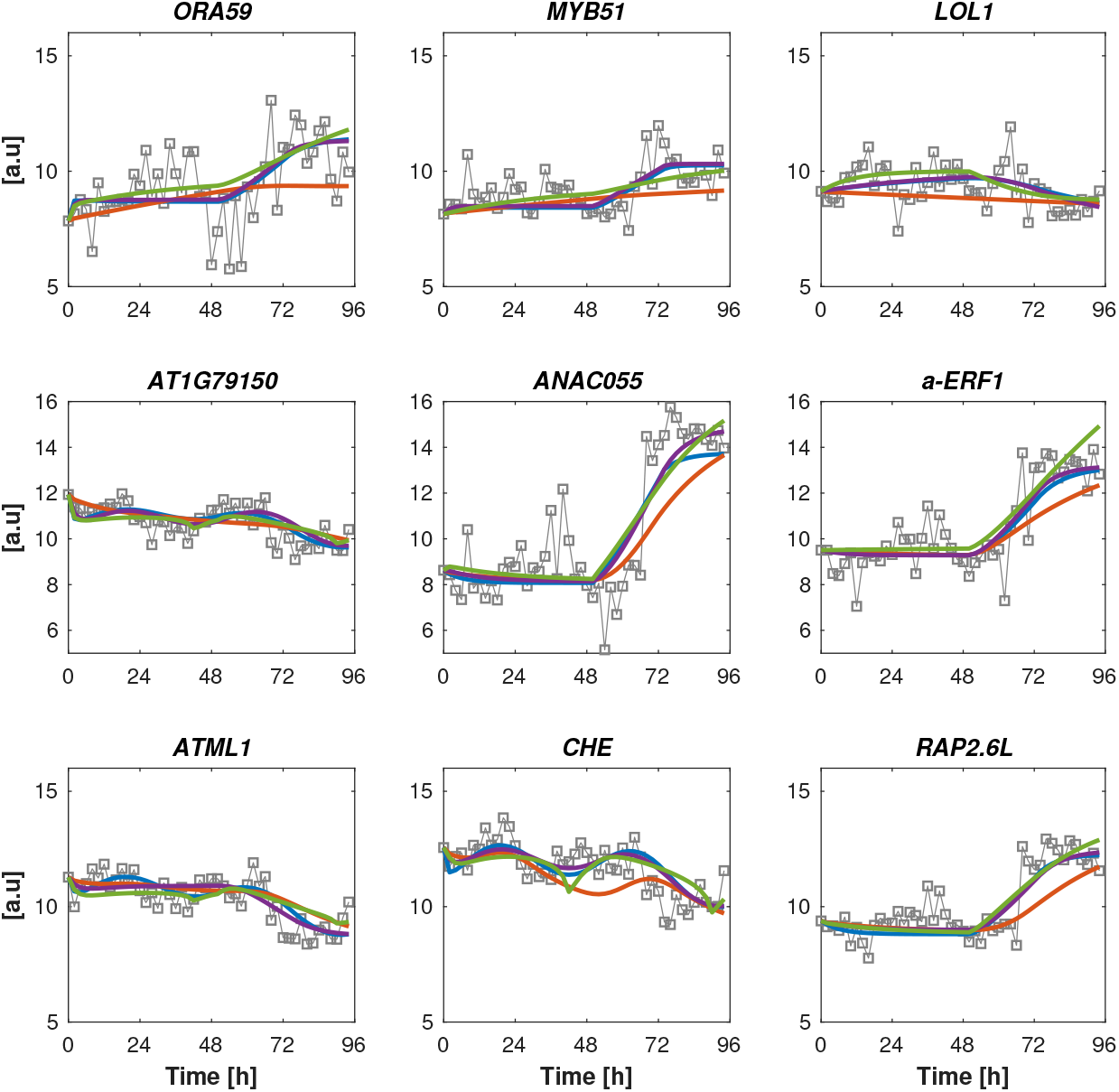
Comparison of the models against experimental data set that is not used in the parameter estimation exercise. Solid grey with ‘square’: Experimental data. Solid blue: Linear model. Solid red: Michaelis-Menten model. Solid green: Standard S-System model. Solid purple: Extended S-System model.

## III. COMPARATIVE ANALYSIS OF THE 9GRN MODELS

### A. Comparison criteria

In this comparative study, the four dynamical models of 9GRN will be evaluated across the following criteria.

- Criterion I: Ease of identifying regulation type.
- Criterion II: Predictive capability.
- Criterion III: Quality of data fit using Akaike weights based on Akaike Information Criterion (AIC).
- Criterion IV: Robustness to parameter uncertainties.

### B. Model structures for 9GRN

The general structure for all these four models are given as follows:

#### Linear model

This linear model is the one used in [6].

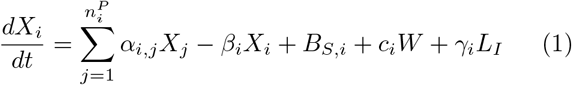

where *X_i_* is the expression level of *i*th gene, 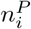 is the number of genes involved in regulating *X_i_, α* is the production rate, *β* is the degradation rate, *B_S_* is the gene basal level while *c* and *γ* parameterised the external input from pathogen *W* and light *L_I_*, respectively. For more details on how each of the terms in (1) are derived, see [6].

#### Standard S-System model

The standard S-System model developed from the field of biochemical system theory was initially proposed in [26] to model metabolic pathways. Over the course of its development (see *e.g.* [27, 28] and references therein), this model has been used to model GRN with good accuracy [29] and it has the following form.

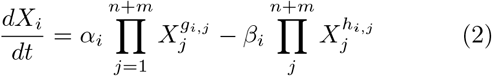

where *α* is the production rate, *β* is the degradation rate, *n* and *m* are respectively the total number of dependent and independent variables and *g_i,j_* and *h_i,j_* are exponents associated with the production and degradation processes, respectively. Note that the standard S-System model structure does not have provision to account for gene basal level and the external input, and these variables are incorporated directly as part of the independent variables.

#### Extended S-System model

This model was proposed in [25] to individually account for the effect of gene basal expression and external input, instead of being part of the independent variables, and was shown to accurately describe the plant circadian system compared to the standard S-System model. The extended S-System model has the following form.

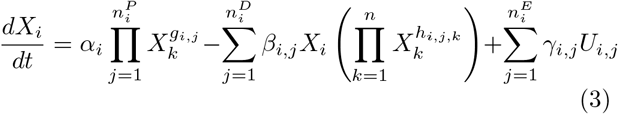

where *α, β* and *γ* are the reaction rate constants associated with production, degradation and external input regulation (*e.g* light, perturbation, basal level, etc), respectively. Like the standard S-System, *g_i,j_* represents the exponent related to production while *h_i,j_* represents the exponent related to degradation. 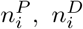 and 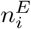 are the number of genetic component involved in the respective production, degradation and external input regulation of *X_i_. U_i,j_* encapsulates the effect of those aforementioned external regulations on *X_i_*.

#### Michaelis-Menten model

Conventionally, this model has been widely used to model GRN (see *e.g.* [30, 31] and references therein) and it has the following form.

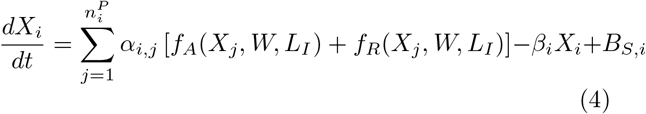

where like before, *α* and *β* are the production and degradation rate, respectively, *B_S_* is the basal level, *f_A_* and *f_R_* are respectively, the activator and inhibitor type of regulation. Both of them have different forms and they are usually modelled as *f_A_* = *X^q^*/(*K^q^* + *X^q^*) and *f_R_* = 1/(*K^q^* + *X^q^*), where *K* is the Michaelis-Menten kinetic constant and *q* is the Hill coefficient. Note that here, the regulations are modelled as a summation of successive regulations and they could also be modelled as a product of successive regulations.

One immediate observation from these four model structures is that the Michaelis-Menten model structure requires the knowledge of the regulation type when deriving the ordinary differential equations (ODE) for each gene, thus making this model not suitable for reverse engineering [32]. If the regulation type is unknown, extra steps (discussed in Section III C) are required to construct the best fitting Michaelis-Menten model structure. Since the Michaelis-Menten model requires extra steps in identifying the regulation types, it can often incur additional computational load.

On the other hand, for the linear and the two S-System based models, the sign of the production rate *α_i,j_* (for linear model) and the exponent associated with the production rate *g_i,j_* (for S-System based model) estimated from data can directly inform us the type of regulation for each gene, where a positive value denotes activation while a negative value denotes inhibition. For the S-System models, there are also approaches being developed that can be used to estimate those parameters in an efficient and fast manner [33]. Moreover, for the linear and extended S-System models, the positive or negative regulation of the external inputs can be also be inferred through the sign of the estimated parameters (i.e., *c*’s and *γ*’s, respectively).

### C. Detailed ordinary differential equations (ODEs) model of 9GRN

We use subscripts *L, SS, ES*, and *MM* in the model parameters to represent the linear, standard S-System, extended S-System and Michaelis-Menten models, respectively. In order to avoid overloading of variables, the following numbers are used to denote the genes in 9GRN. 1: *ORA59,* 2: *MYB51,* 3: *LOL1,* 4: *AT1G79150,* 5: *ANAC055,* 6: *a-ERF-1,* 7: *ATML1,* 8: *CHE* and 9: *RAP2.6L*.

#### Linear model

The corresponding ODEs following (1) are given as follow, which is the same linear model used in [6],

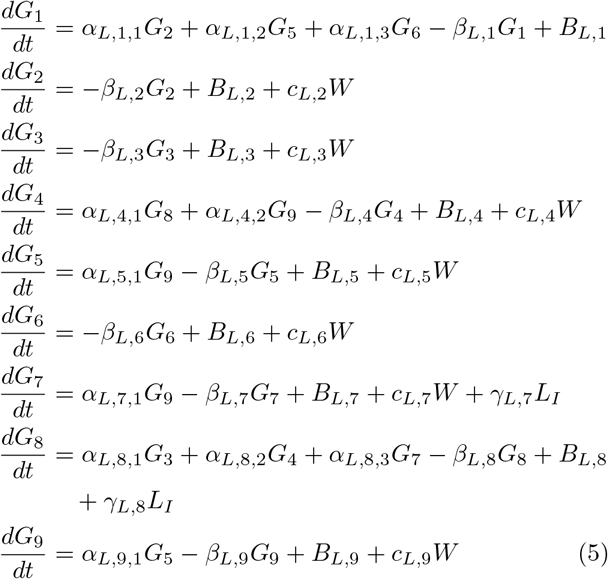

#### Standard S-System model

Following (2), the corresponding ODEs for 9GRN are given as follow.

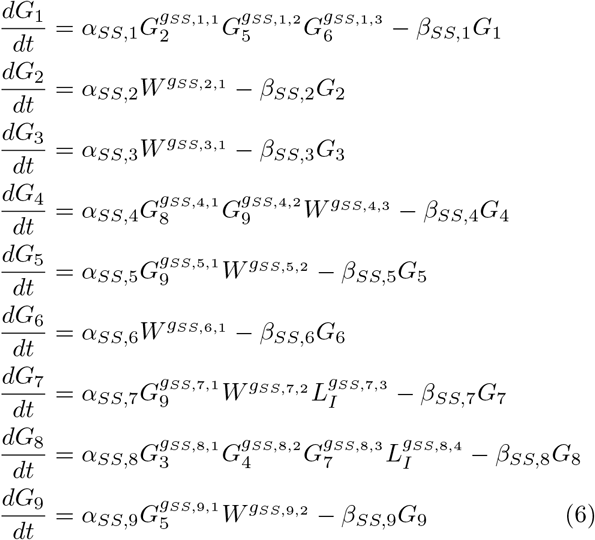

Note again that the two external variables *W*, which represents the effect of *Botrytis cinerea* inoculation and *L_I_*, which represents the effect of light are considered as the independent variables.

#### Extended S-System model

Following (3), we arrive at the following ODEs for the 9GRN,

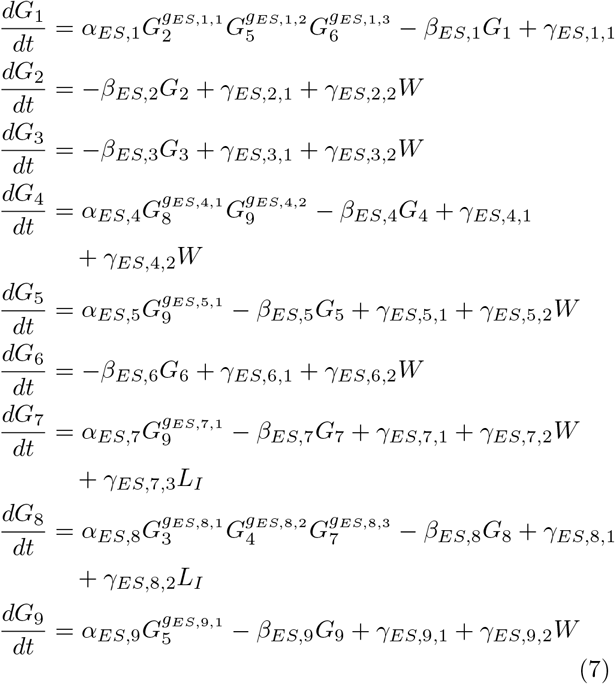

As a remark, despite the regulation type is unknown, the ODEs for these three models can still be written down as depicted in (5), (6) and (7), as the regulation type can be inferred through the sign of the estimated parameters. Also, for the S-System based models, we set *h_i,j_* = 1 to reduce the amount of parameters that need to be estimated.

#### Michaelis-Menten model

Unlike the previous three models, the ODEs of the Michaelis-Menten model can only be written down when the type of regulation is known. To facilitate the derivation of these ODEs, we need to employ additional steps to infer those regulation types.

GRN network inference and parameter estimation using Michaelis-Menten ODEs can be a challenging problem, as repeatedly solving the ODEs via numerical integration can be computationally expensive. In this study, we employ our recently proposed parametric gradient-matching method (see Supplementary Text Section S1.3, Algorithm I and [34]) as the GRN inference approach, which incorporates dynamics information and computational efficient. It is an inference approach based on parametric Michaelis-Menten nonlinear ODEs representation of a GRN [35]. The approach significantly reduced the computational cost of repeatedly solving the candidate ODEs via a two-step gradient matching. It first employs a Gaussian process to interpolate each time-course gene expression data. Then, the parameters of the ODEs are optimised by minimising the difference between interpolated derivatives and the right-hand-side of the ODEs. In such a way, the ODEs do not need to be solved explicitly, thereby reducing the computational cost. For more details of the method, see [34, 35]. We note that there are copious of other similar methods to identify regulation type of the Michaelis-Menten model in a GRN (see *e.g.* [36, 37]). As the main goal of this work is to perform comparative analysis of the GRN models and not on the network inference algorithm, we will treat the identified regulation type from our network inference algorithm as the correct regulation for our comparative analyses. The summary of the identified regulation types is given in Table I.

With that, the corresponding ODEs are given as follow.

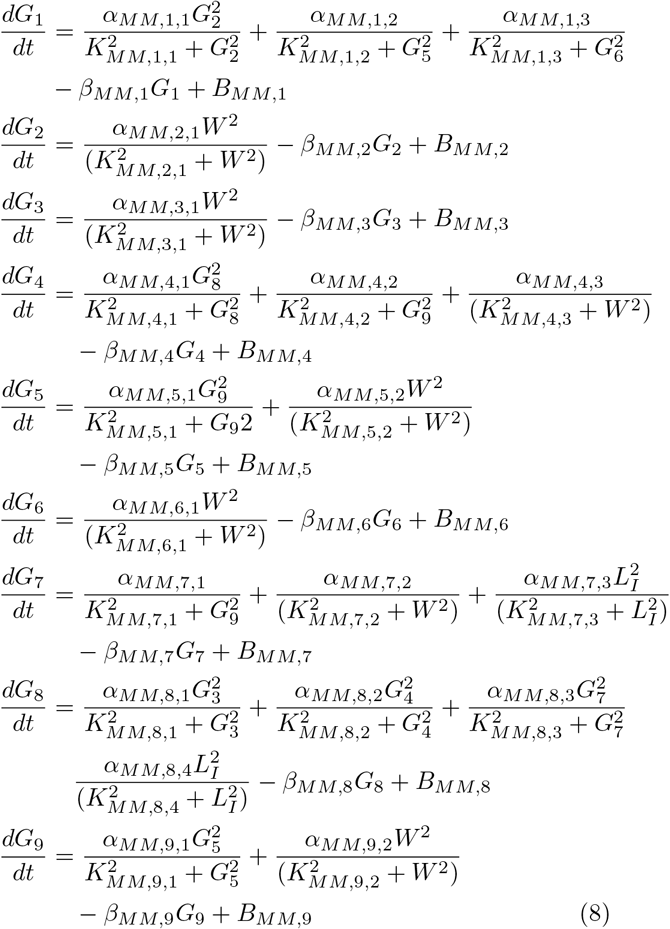

**TABLE I:** Identified regulation types for interaction within 9GRN following the parametric gradient-matching approach. The (+) and (-) signs indicate the activation and inhibition regulation types respectively. The signs for *W* and *L_I_* indicate that this gene is positively or negatively regulated by those external inputs (see Fig. 1).

| Number | Gene | Regulation Types |
| --- | --- | --- |
| 1 | *ORA59* | *MYB51* (+), *ANAC055* (-), *a-ERF-1* (-) |
| 2 | *MYB51* | $W$ (+) |
| 3 | *LOL1* | $W$ (+) |
| 4 | *AT1G79150* | *CHE* (+), *RAP2.6L* (-), $W$ (-) |
| 5 | *ANAC055* | *RAP2.6L* (+), $W$ (+) |
| 6 | *a-ERF-1* | $W$ (+) |
| 7 | *ATML1* | *RAP2.6L* (-), $W$ (-), $L_I$ (+) |
| 8 | *CHE* | *LOL1* (+), *AT1G79150* (+), *ATML1* (+), $L_I$ (+) |
| 9 | *RAP2.6L* | *ANAC055* (+), $W$ (+) |

In all the four models, the infection of *Botrytis cinerea* is modelled as a step function with gradual increase from time 48 to 72 hours, *i.e.* the time inoculation occurs. Mathematically, this is modelled as

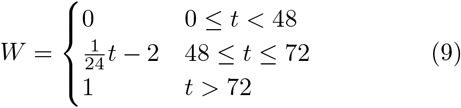

For the light regulated genes, these genes are affected by the duration of photoperiod of light. In [10], the experiment was carried out under 16 hours of light and 8 hours of dark. The resulting genes in response to this photoperiod duration behave in a sinusoidal manner with its peak between 8 to 10 hours at the first instance of light. In view of this, the effect of light is modelled as a sinusoidal signal that peaks at around 9 hours at the first stance of light and its expression is given by

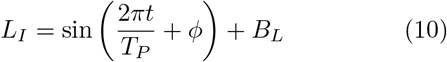

where *B_L_* = 1.0001 is the expression base level, *ϕ* = *π*/6 radian is the phase shift and *T_P_* = 24 hours is the period of the sinusoid. The reason for setting *B_L_* = 1.0001 is to avoid *L_I_* becoming zero, which can be problematic when it is used in modelling 9GRN using standard S-System.

### D. Parameter estimation

The data used in this study is taken from [6]. For the linear and two S-System based models, the parameters of the corresponding models were fitted to the experimental data set by minimising the weighted mean squared error (WMSE) between the simulated and experimental data, *i.e.* by finding

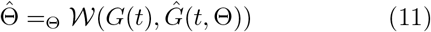

where

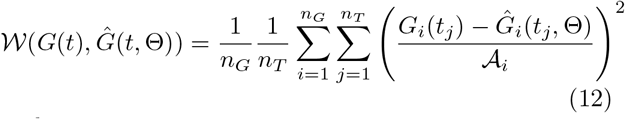

with

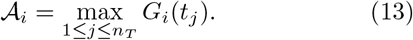

where *G* represents the gene component, *t* denotes the time index, *n_G_* = 9 is the number of gene components and *n_T_* = 48 is the number of data point used. Given that the amplitude of different gene components is different, to allay any bias during the optimisation procedure for model parameter fitting, we introduce the weights 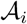 in (12) where we normalise each time series to its maximum value. The MATLAB function `fminsearch` that employs Nelder-Mead simplex algorithm was used to minimise (11). The estimated parameters for these three models are given in Tables S1 to S3. Note that the estimated model parameters of the linear model is somewhat different than the one provided in [6]. This is because in this study, instead of directly using the estimated parameters from [6], they are used as the initial value for the optimisation to determine whether any further improvement in terms of the WMSE can be achieved. The estimated parameters given in Table S1 are very close to the one estimated in [6] suggesting a high confidence level in the estimated parameters for the linear model.

The parameters associated with the Michaelis-Menten model are given in Table S4. These parameters have been estimated together with the inference algorithm (see Supplementary Text, Section S1.3) via the gradient-matching method (see [34] and its Supplementary Material for more details).

The identified regulation types for these three models are given in Table II. We have also included the regulation types inferred from Michaelis-Menten model in this table for ease of comparison. In general, there is a general consensus on the identified regulation types shown in Table II apart from genes *ORA59, LOL1* and *AT1G79150*. Specifically, for gene *ORA59,* only the inferred regulation type for *a-ERF-1* when using the Michaelis-Menten model is different from the other three models. For *ORA59*, the time series shows an increasing trend, which is consistent with the increasing trend of *a-ERF-1*, suggesting a higher possibility of a positive regulation, which agrees with the three models rather than the Michaelis-Menten model. For gene *LOL1*, there is difference in the inferred pathogen regulation type with the linear and standard S-System models identified negative regulation, while extended S-System and Michaelis-Menten models identified positive regulation. Lastly, for gene *AT1G79150,* the inferred regulation types for *RAP2.6L* and pathogen are different across all four models. A detail look at the time series data for these genes *LOL1* and *AT1G79150* (Fig. S1) suggests that the difficulty in identifying these regulation types is attributed to the almost plateau nature of these two gene expression levels.

**TABLE II:**
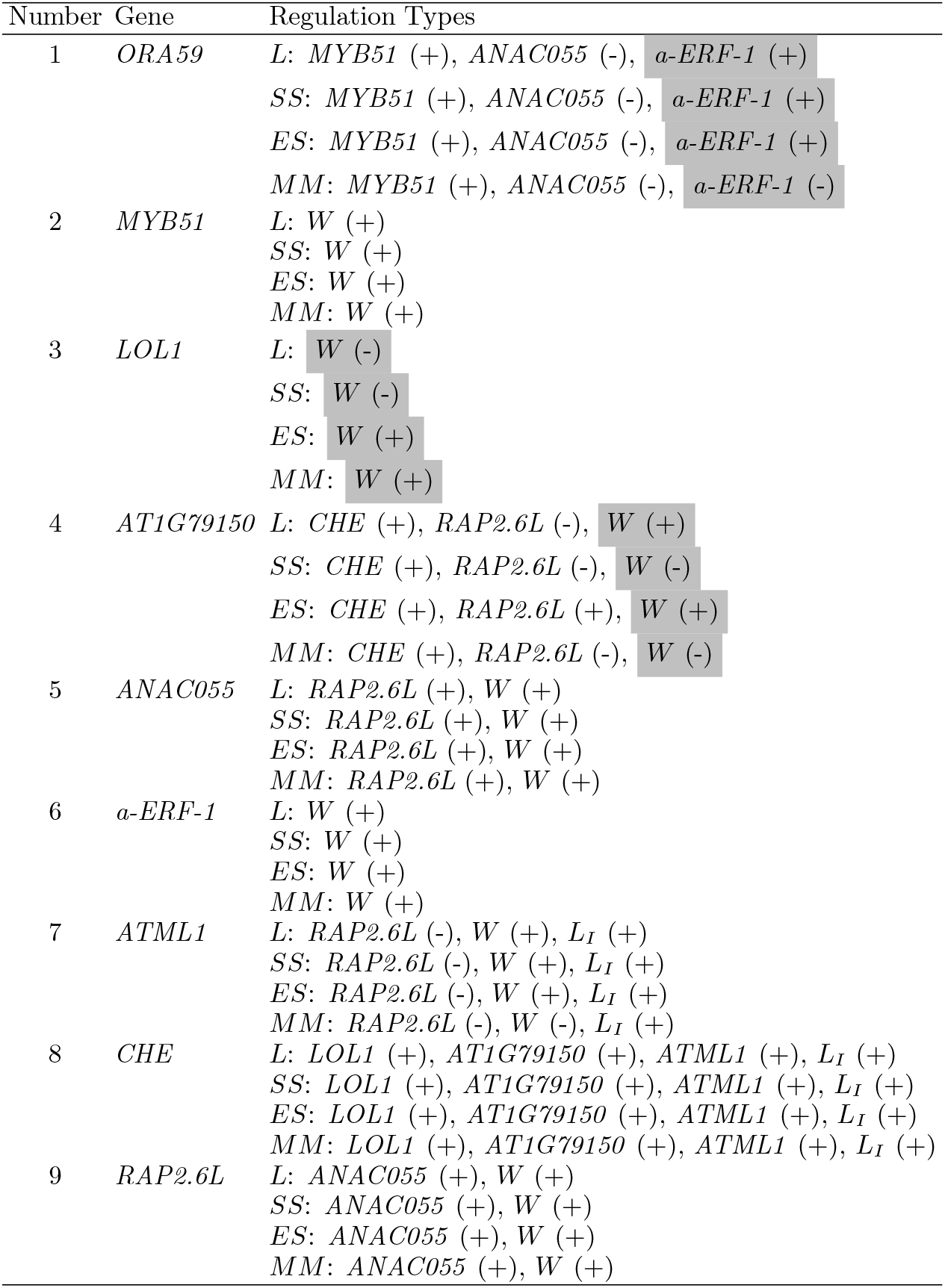
Identified regulation types based on estimated parameters of 9GRN using linear (L), standard S-System (*SS*) and extended S-System models (ES). The (+) and (-) signs indicate the activation and inhibition regulation types, respectively. The signs for *W* and *L_I_* indicates that this gene is positive/negative regulated by those external inputs (see Fig. 1). The regulation type for Michaelis-Menten model (*MM*) shown in Table I is also listed for ease of comparison. Identified regulation types that are different are highlighted in grey.

Using the identified parameters given in Tables S1 to S4, we compared the predictive capability of the models with the experimental data on a set of data that is not used in parameter estimation exercise and the result are shown in Fig. S1. As a quantitative measure, we calculated the WMSE, using (12), and they are shown in TABLE III.

**TABLE III:**
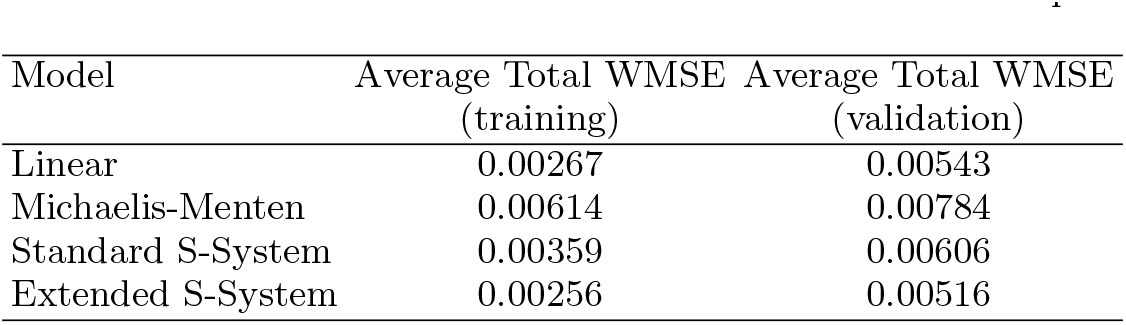
Average total WMSE for both ‘training’ and ‘validation’ data sets for 9GRN, which is calculated by taking the average sum of the individual WMSE given in Tables S5 and S6. The ‘training’ data set refers to the data that is used in parameter estimation exercise, while the ‘validation’ data set refers to the data that is not used in the parameter exercise.

The results shown in Fig. S1 and Table III shows that the all four models are able to pick up the general trend of the data well. Specifically, the linear and two S-System models perform really well with relatively smaller total WMSE compared to the Michaelis-Menten model. For the Michaelis-Menten model, there are several instances where the model fall short in terms of realising the correct amplitude levels (e.g. *ORA59* and *MYB51*), which is also reflected in the individual WMSE shown in Tables S5 and S6.

To further test the performance of these models, we compare qualitatively the dynamics of these four models against mutant data set, where two different genes, *i.e.*, Δ*nac* and Δ*rap*2.6*l* have been mutated. Fig. 3 shows the predictive capability of the four models against the mutant data. In general, all models pick up the correct trend of the mutant behaviours albeit the two S-System models have difference in the amplitude. For instance, gene *ORA59* from the extended S-System model has higher expression level under both knockdown mutants. Similarly, gene *RAP2.6L* from standard S-System model has lower expression level under Δ*anac*055. Nevertheless, all the models are able to predict the mutant behaviours qualitatively well. Readers who are interested in the quantitative mutant analysis can refer to Supplementary Text.

**FIG. 3:**
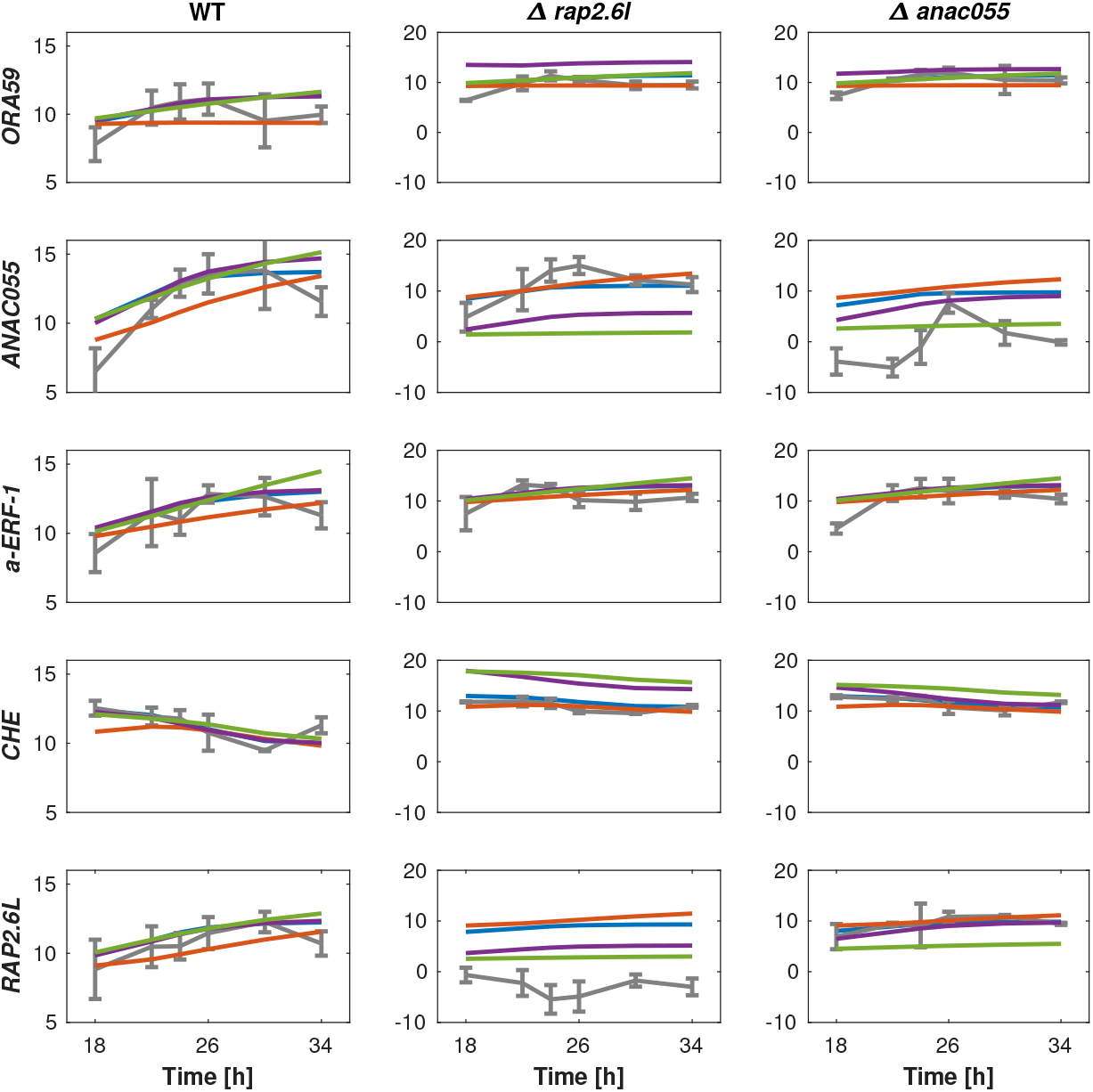
Comparison of the models against mutant experimental data set. For the simulation of the mutant, we reduce the production rate associated with the knockdown gene by 20%. Solid grey with error bar: Experimental data. Solid blue: Linear model. Solid red: Michaelis-Menten model. Solid green: Standard S-System model. Solid purple: Extended S-System model.

### E. Assessing model quality using Akaike weight based on Akaike Information Criterion (AIC)

While the WMSE and the mutant analysis provide respectively the quantitative and qualitative approaches of the performance of the model, these approaches however do not reflect fully the quality of fit given the different model structures employed and the number of parameters used. In order to quantify the relative quality of the model fits to the experimental training data obtained with the four models considered, we employed the widely-used Akaike Information Criterion (AIC), which calculates the best approximating model to a given dataset with respect to Kullback-Leibler information loss [38, 39].

For a given model, the AIC is defined as

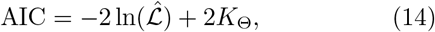

where 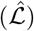 is the maximised log-likelihood and *K*_Θ_ is the number of model parameters. Consider that the optimal parameter estimates for all four 9GRN models were acquired through the minimisation of weighted least squares cost function, it can be shown that [40]

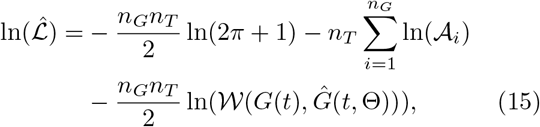

with *n_G_* is the number of genes, *n_T_* is the number of data points in the time series, 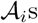 as defined in (13) are the cost function weights and 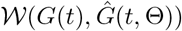 as defined in (12).

By denoting AIC_*i*_ as the AIC value of the *i*th model, these four 9GRN models are ranked by their AIC differences calculation, *i.e.*

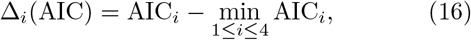

and finally, the corresponding Akaike weights can be calculated as follow:

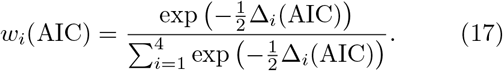

We can interpret this Akaike weight, *w_i_*(AIC) as the probability that the ith model is the best from the perspective of minimising K-L information loss, given the set of candidate models and the data. In addition, the strength of evidence that favours model *i* over model *j* is quantified by the ratio *w_i_*(AIC)/*w_j_* (AIC) [38–41].

Finally, since *n_G_, n_T_* and 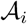 in (15) are fixed across the respective GRN models, the expression of the AIC value (*i.e.,* (14)) can be further simplified to

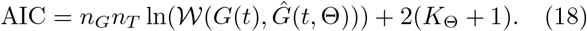

where (18) is then used to compute the AIC differences Δ_*i*_(AIC) and Akaike weights *w_i_*(AIC) of a given 9GRN model.

The AIC criterion in Table S9 indicates that the two most viable candidate models (in the sense of K-L divergence) are the extended S-System model and the linear model with their Akaike weight of *w_ES_*(AIC) = 0.9836 and *w_L_*(AIC) = 0.0164, respectively. Between these two models, the ratio of *w_ES_*(AIC)/*w_L_*(AIC) ≈ 60 suggests that the extended S-System model is 60 times more likely to be the viable model candidate compared to the linear model. On the other hand, the Akaike weights also exclude the Michaelis-Menten and the standard S-System models as the viable models given their Akaike weights are close to zero.

### F. Robustness of the models to parameter uncertainties

In practice, the estimated parameters of the model are subjected to uncertainties (e.g. intrinsic noise, modelling error, etc). To test the robustness of these four models, we perform a global sensitivity analysis, where all the parameters of the model are simultaneously varied in a random manner in each simulation, In this study, we assume that the uncertainties account for the parameters to vary ±30% (see *e.g.* [42–44]) from its nominal value. To ensure an unbiased sampling of the parameter values, we adopted the *Latin Hypercube Sampling* approach (see *e.g.* [45, 46]) to randomly generate a parameter set that is within ±30% of the original value of each parameter for each simulation.

In the Latin Hypercube Sampling approach, each model parameter is first discretised into *N_s_* evenly spaced intervals from the defined lower and upper bounds. As we are varying the parameter within ±30%, this results in *N_s_* evenly spaced interval between 0.7× to 1.3× the nominal parameter. Here, we choose *N_s_* = 1000, and this results in a total number of (1000 × Kθ) randomly combined parameter sets, where Kθ is the number of parameter in each of the four models. We run a total number of 10000 simulations for each of the four models, where in each simulation, we sample only once from this total number of randomly combined parameter sets. Due to the non-repetitive nature of this sampling approach, not only the biased sampling can be averted, an extensive sampling within the model parameter range of interest can also be covered [45].

Following [22], we compute the Mean Relative Error (MRE) given by

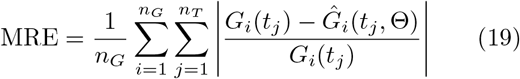

as a quantitative metric to evaluate the model response to parameter uncertainties using the same notation as (12). To determine the robustness of the models, we collate the number of simulations (over 10000), where the MREs are within 4× the nominal MRE value. The choice of 4× is based on the observation over 10000 simulations that the performance of the models are deemed acceptable. To compare the robustness of the model, a model is considered more robust than the other if the number of simulations within 4× nominal MRE value is higher in the former than the latter.

Table V shows the MRE values for all four models. Defining *N_SIM_* as the number of simulation for each model where the MRE values are within 4× nominal MRE value. The results show that the Michaelis-Menten and the extended S-System models has the largest and smallest *N_SIM_* values, respectively suggesting these models respectively being relatively the most and least robust to parameter uncertainty. Also, we notice that the *N_SIM_* values for the two S-System based models are comparative smaller, which is expected given that the exponent term tends to be sensitive to uncertainties [47].

**TABLE IV:** Ranking model fits to experimental data based on AIC weights for 9GRN. The notation *L, MM*, *SS, ES,* denote the linear, Michaelis-Menten, Standard S-System and Extended S-System models, respectively. Here *n_G_* = 9, *n_T_* = 48, *K*_Θ_ is the number of parameters in the model, 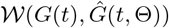 is the WMSE best fit to the data set used for parameter estimation, Δ_*i*_(AIC) is the AIC differences and *w_i_*(AIC) is the Akaike weights for each model.

| Model | $L$ | $MM$ | $SS$ | $ES$ |
| --- | --- | --- | --- | --- |
| $K_\Theta$ | 38 | 58 | 38 | 44 |
| $\mathcal{W}(G(t), \hat{G}(t, \Theta))$ | 0.00267 | 0.00614 | 0.00359 | 0.00256 |
| $\Delta_i(\text{AIC})$ | 8.182 | 397.506 | 134.696 | 0 |
| $w_i(\text{AIC})$ | 0.0164 | $4.14 \times 10^{-89}$ | $5.55 \times 10^{-30}$ | 0.9836 |

**TABLE V:** Nominal MRE for each model and the number of simulation across 10000 that has the MRE values within 4× the nominal MRE value. The notation *L, MM, SS, ES*, denote the linear, Michaelis-Menten, Standard S-System and Extended S-System models, respectively. *N_SIM_* denotes the number of MRE within 4× nominal MRE value

| Model | $L$ | $MM$ | $SS$ | $ES$ |
| --- | --- | --- | --- | --- |
| Nominal MRE | 0.0523 | 0.0704 | 0.0516 | 0.0611 |
| $N_{SIM}$ | 2285 | 9271 | 1082 | 731 |

To see how the *N_SIM_* values are distributed, we plot the histogram in Fig. S2, and the histogram shows that the majority of the MRE values are distributed close to the nominal MRE value indicating these models are more robust than anticipated. To further investigate this, we plot the lower and upper bound of each model simulated using the parameter sets within *N_SIM_* that produce the largest and smallest MRE value and these plots are shown in Figs. S3 to S6. Interestingly, majority of the genes are robust to parameter uncertainty where their uncertainty bounds are narrow apart from a handful of genes (*e.g. ORA59, ANAC055* and *CHE*), where we observe a wider uncertainty bounds. Moreover, despite having the largest *N_SIM_* value, the Michaelis-Menten model has four genes with wide uncertainty bounds compared to the same four genes for the other three models. This suggests that while the larger *N_SIM_* in Michaelis-Menten model is most probably attributed to the genes with narrow uncertainty bound, it comes at the expense of reduced robustness in other genes such as *CHE*, which has the widest uncertainty bound.

## IV. DISCUSSION AND CONCLUSION

In this study, we have compared four dynamical models of 9GRN obtained using a data-driven modelling approach in terms of four criteria, namely their ease of identifying regulation type, predictive capability, quality of data fit based on AIC and robustness to parameter uncertainties.

The linear and the two S-System based models have a general model structure that can facilitate the identification of the regulation types directly from data through the sign of the estimated parameters. In contrast, due to the requirement of different functions for different regulation types for the Michaelis-Menten model (Section III B), additional steps are required to ensure the most viable regulation types when identifying them from data, making this model the least favoured in terms of Criterion I. Furthermore, despite the identified regulation types given in Table II showing a consensus, when comparing the difference in the identified regulation types, the linear and two S-System based models have more common agreement compared to the Michaelis-Menten model for *e.g.* in gene *ORA59.*

In terms of the model predictive capability, the linear and extended S-System models rank higher in terms of their smaller WMSE value both in the training and validation data set compared to the standard System and Michaelis-Menten models (Table III) suggesting Criterion II is in favour of these two models. In terms of mutant analysis, between the linear and extended S-System models, the former qualitatively better predicts the mutant behaviours (Fig. 3) than the latter.

For Criterion III, the analysis of AIC weights (Table S9) suggests the linear and extended S-System models are the two most viable candidate models compared to the standard S-System and Michaelis-Menten models. Nevertheless, the extended S-System model is 60 times more likely to be the candidate model compared to the linear model given its larger AIC weight, *w_ES_*(AIC), which suggests the extended S-System in the most favoured model for Criterion III.

For the last criterion, the analyses using Latin Hypercube Sampling and MRE (Table V) indicate that the Michaelis-Menten model has the largest *N_SIM_*, suggesting this model is relatively robust against parameter uncertainty compared to the other three models. Interestingly, when analysing the histogram of the MRE distributions (Fig. S2) and the lower and upper bounds uncertainty plots (Figs. S3-S6), the width of the uncertainty bounds are smaller and similar across the linear and the two S-System based models. On the other hand, despite the Michaelis-Menten model having narrow uncertainty bounds across most of the genes in 9GRN, some genes (e.g. *CHE* and *ORA59*) have the widest uncertainty bound across all four models. This suggests that the large *N_SIM_* of the Michaelis-Menten model are attributed to the narrow bounds of most genes but at the expense of wide bounds on certain genes like *CHE*.

Table VI summarises the performance of all four models across the four criteria. For Criteria II to IV, we provide the associated ranking in each criterion with ‘1’ being the most favoured model and ‘4’ being the least favoured model based on the metrics used to compare them. We then calculated the Total Rank Score (TRS), which is the sum of the ranking number given in bracket with the smallest and largest scores represent the most and least favoured models, respectively.

**TABLE VI:** Summary of the model performance across four criteria. For Criteria II to IV, the model is ranked in bracket with ‘1’ being the most favoured model and ‘4’ being the least favoured model according to the metric used in the comparison. The notation *L, MM, SS, ES*, denote the linear, Michaelis-Menten, Standard S-System and Extended S-System models, respectively. For Criterion II, the notation ‘Tra.’ and ‘Val.’ represent training and validation, respectively. The Total Rank Score (TRS) is the sum of the ranking number across Criteria II to IV given in the bracket.

| Criterion | $L$ | $MM$ | $SS$ | $ES$ |
| --- | --- | --- | --- | --- |
| I | Easy due to its general model structure | Difficult due to extra steps required to determine the relevant function | Easy due to its general model structure | Easy due to its general model structure |
| II | Tra. WMSE = 0.00267 (2)<br>Val. WMSE = 0.00543 (2) | Tra. WMSE = 0.00601 (4)<br>Val. WMSE = 0.00768 (4) | Tra. WMSE = 0.00359 (3)<br>Val. WMSE = 0.00606 (3) | Tra. WMSE = 0.00256 (1)<br>Val. WMSE = 0.00516 (1) |
| III | $w_L(\text{AIC}) = 0.0164$ (2) | $w_{MM}(\text{AIC}) \approx 0$ (4) | $w_{SS}(\text{AIC}) \approx 0$ (3) | $w_{ES}(\text{AIC}) = 0.9836$ (1) |
| IV | $N_{SIM} = 2285$ (2) | $N_{SIM} = 9271$ (1) | $N_{SIM} = 1082$ (3) | $N_{SIM} = 731$ (4) |
| TRS | 8 | 13 | 12 | 7 |

The extended S-System model scores the smallest TRS, followed closely by the linear model, while the Michaelis-Menten model scores the largest TRS. While the linear model scores a lower TRS compared to the extended S-System model, the linear model performs consistently across all criteria with rankings of ‘2’ compared to the extended S-System model. Based on this consistency, we surmise that the linear model is a more viable candidate model for constructing this 9GRN using a data-driven modelling approach.

The finding from our comparative analysis is somewhat consistent with the finding from [22], where in that study, the standard S-System model is found to be a more viable model compared to the Michaelis-Menten and massaction model for describing plant flowering time regulatory network. Between the standard and extended S-System models in our study, our analysis shows that the latter model outperforms the former model across the given criteria. This is expected given that the extended S-System model considers the external input as being a separate term instead of grouping them as part of the dependent variables. This thus provides more degree of freedom for the external input to influence the model dynamics, which could improve the accuracy of the model [25].

Our finding that the least viable model being the Michaelis-Menten model may seem surprising given its wide usage in modelling GRN. In a review work by [48], it has been reported that the Michaelis-Menten rate law has been often misused without ensuring the valid operating condition in many previous studies. The same review (and references therein) and our previous studies [25, 49] also highlighted issues pertaining to the identifiability of the Michaelis-Menten parameters. These two points accentuated the underlying challenge in using Michaelis-Menten model, which could possibly be the reason for its poor viability. One may potentially argue that the choice of network inference algorithm to obtain the Michaelis-Menten models (such as the one used in this study) may influence the analysis and the results. As such, we derive an alternate Michaelis-Menten model with the regulation type following the linear model and found that despite showing some improvement in Criteria II and III, the overall performance of the model is still ranked behind the linear and extended S-System model (see Tables S8 and S9).

Returning to our main question posed for this study — “*In using the data-driven modelling approach, what is the most viable model given the temporal data and knowledge about the 9GRN interaction*”? While traditionally Michaelis-Menten model has been the model of choice due to its biological relevance (see *e.g.* [32]), our comparative analysis seems to tip the balance towards the linear model being the preferred choice of model for 9GRN suggesting the linear model used for genetic control design suggested in [6] is a viable one. Our results also suggest that both linear and the extended S-System models are good alternatives for modelling GRN as compared to the commonly used Michaelis-Menten model, which is in agreement with previous studies (see *e.g.* [22, 29]).

## AVAILABILITY

All the MATLAB simulation codes are available at https://github.com/mathiasfoo/9grncomparison

## AUTHORS CONTRIBUTION

MF and FH conceived the study. MF and LD performed the simulation and analysed the data. MF and FH drafted the manuscript. All authors read and approved the manuscript.

## ACKNOWLEDGEMENTS

MF would like to thank Dr. Xun Tang from Louisiana State University for providing assistance with the global sensitivity analysis using Latin Hypercube Sampling.

## S1 SUPPORTING INFORMATION

### S1. EXTENDED METHODS

#### A. Total Mean Square Error Calculation for Mutant Analysis

The mutant analysis given in Figure 3 of the main text shows that all models are able to predict the general trend of the mutant behaviour well in qualitative manner, which is encouraging given that these mutant data independent data set that are not used in parameter estimation exercise.

Nonetheless, we compute the Total Mean Square Error (TMSE) for the mutant analysis to provide some form of quantitative analysis in order to provide model ranking as shown in Table 6 of the main text. Note that the calculation of TMSE is purely for model ranking purpose.

The TMSE can be calculated in the following manner

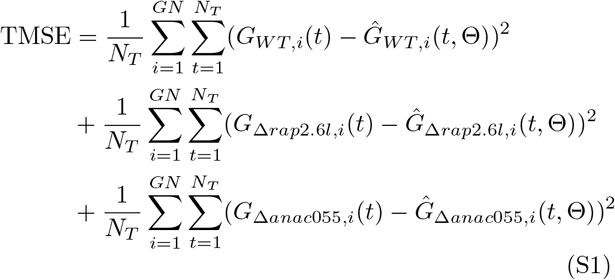

where *GN* ∈ {*ORA59, ANAC055, a-ERF-1, CHE, RAP2.6L*}, *N_T_* is the total number of data point, *G* represents the data set, *Ĝ* represents the predicted data from model and the subscripts *WT*, Δ*rap*2.6*l* and Δ*anac*055 represent the Wild Type, *RAP2.6L* mutant and *ANAC055* mutant, respectively. The calculated TMSE are shown in Table S7.

#### B. Alternate Michaelis-Menten Model

To determine whether the network inference algorithm would affect the performance of the Michaelis-Menten model structure, we consider an alternate Michaelis-Menten model structure where its regulation types follow the linear model. With that, the corresponding ODEs are given below and the estimated parameters following the parameter estimate approach given in main text is given in Table S10.

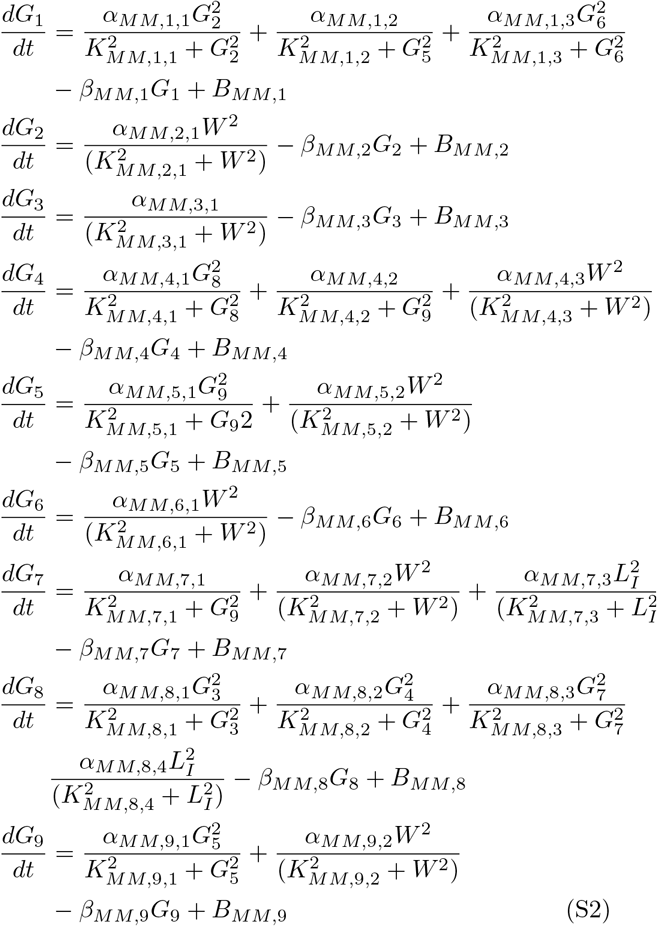

#### C. Network Inference Algorithm for Michaelis-Menten Model Structures

The following summarised algorithm has been used in this study to identify the regulation type of the Michaelis-Menten model structure for 9GRN. For more details, see Dony et al. (2019).

Algorithm I: GRN inference with parametric-gradient matching:

1. Gene expression time-course data interpolation with Gaussian Processes (GP).
2. Gene expression data derivatives computation with GP.
3. Optimise the parameters of all possible ODE models (network topologies) using gradient-matching.
4. Model selection. Compute the edge weighting for each type of regulatory interaction, based on BIC criterion and the likelihood (or distance) of each gene with respect to their possible parents. Therefore, the best network topology and types of regulatory interactions can be determined.
5. Evaluate the overall performance of the GRN inference. The Area Under the Precision-Recall (AUPR) curve is calculated based on the BIC weights of every edge in the network.

### S2. SUPPLEMENTARY FIGURES AND TABLES

**FIG. S1:**
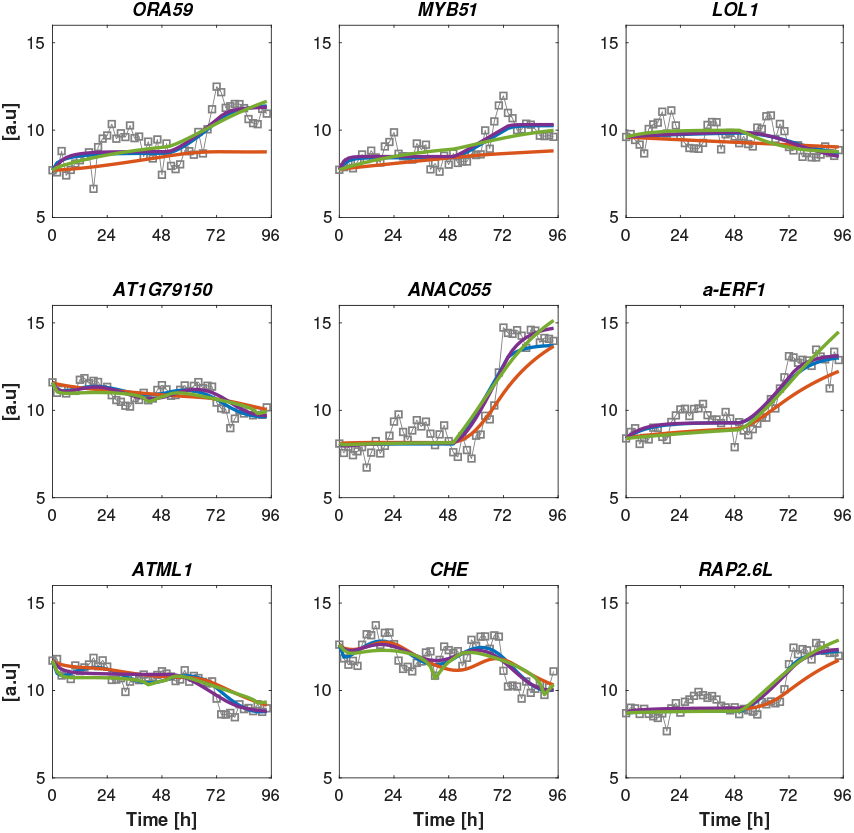
Comparison of the models against experimental data set that was used in the parameter estimation exercise. Solid grey with ’square’: Experimental data. Solid blue: Linear model. Solid red: Michaelis-Menten model. Solid purple: Extended S-System model. Solid green: Standard S-System model.

**FIG. S2:**
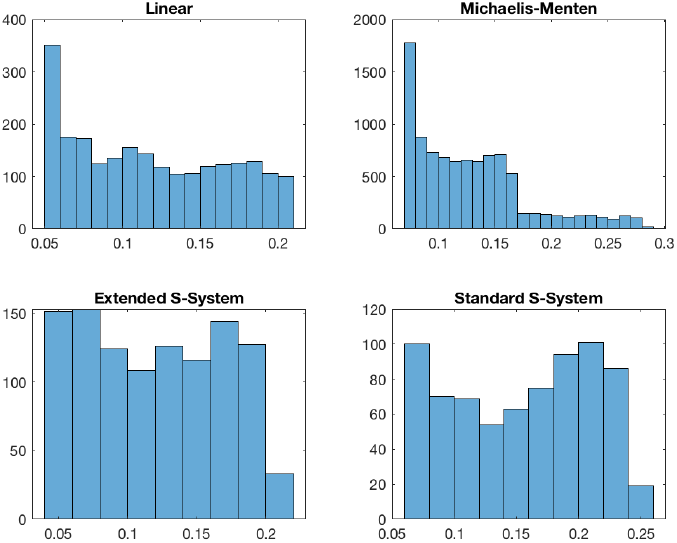
Histogram of the MRE values across 10000 simulations where the MRE values are within 4× the nominal MRE value.

**FIG. S3:**
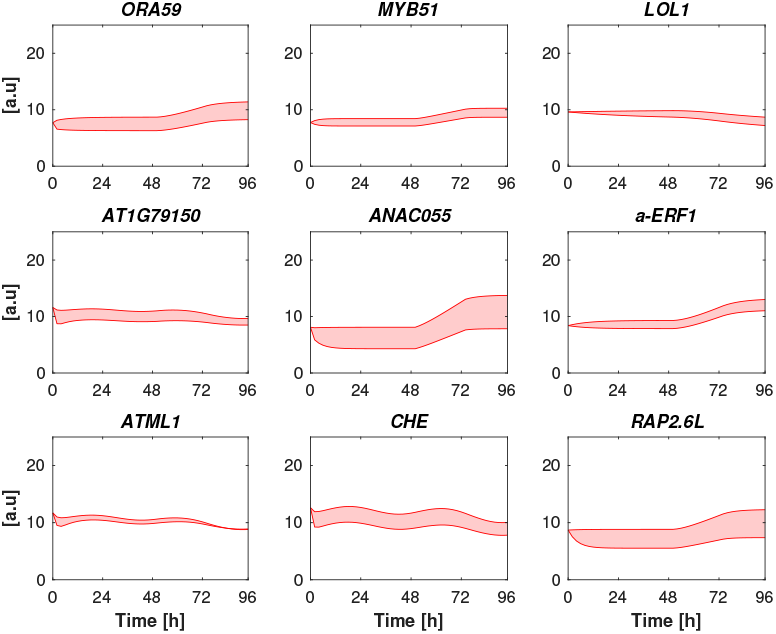
Upper and lower bound of the linear model shown in shaded red that is subjected to parameter uncertainties. The upper and lower bound are taken from the number of simulation where the MRE values are within 4× the nominal MRE value.

**FIG. S4:**
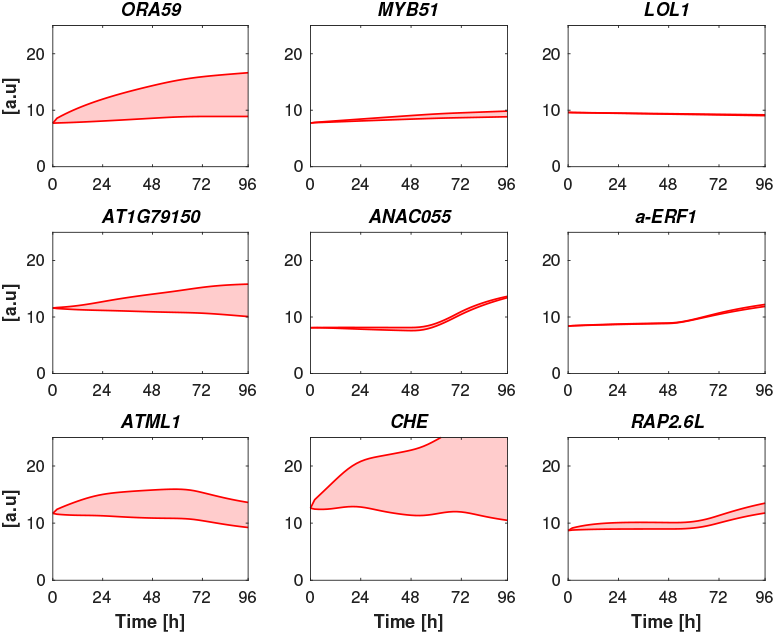
Upper and lower bound of the Michaelis-Menten model shown in shaded red that is subjected to parameter uncertainties. The upper and lower bound are taken from the number of simulation where the MRE values are within 4× the nominal MRE value.

**FIG. S5:**
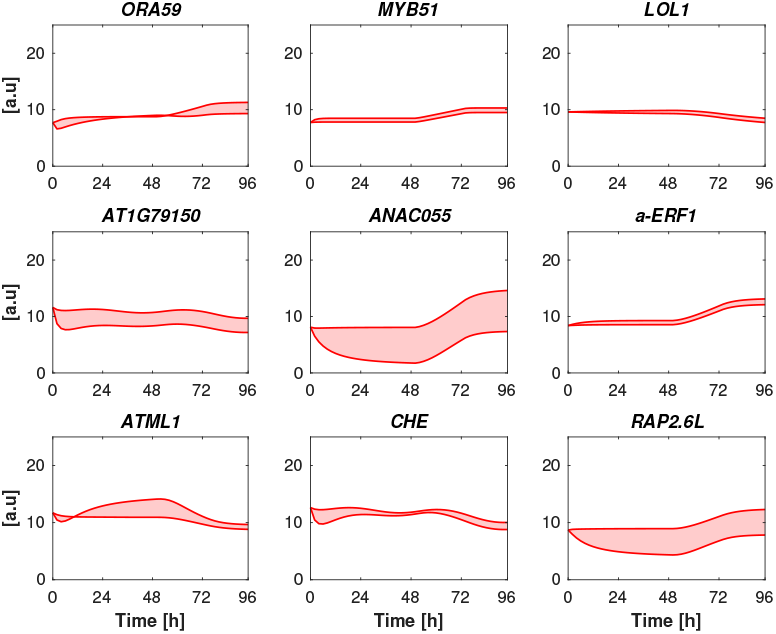
Upper and lower bound of the extended S-System model shown in shaded red that is subjected to parameter uncertainties. The upper and lower bound are taken from the number of simulation where the MRE values are within 4× the nominal MRE value.

**FIG. S6:**
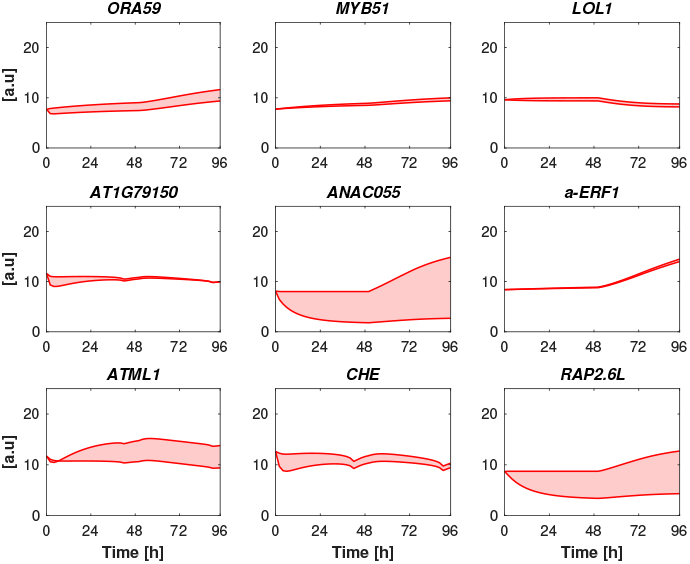
Upper and lower bound of the standard S-System model shown in shaded red that is subjected to parameter uncertainties. The upper and lower bound are taken from the number of simulation where the MRE values are within 4× the nominal MRE value.

**TABLE S1:** Estimated parameters of 9GRN using linear model

| Number | Gene | Parameter |
| --- | --- | --- |
| 1 | *ORA59* | $\alpha_{L,1,1} = 14.4341, \alpha_{L,1,2} = -0.7477, \alpha_{L,1,3} = 23.6810$<br>$\beta_{L,1} = 40.8322, B_{L,1} = 18.8299$ |
| 2 | *MYB51* | $\beta_{L,2} = 0.6765, B_{L,2} = 5.6837, c_{MYB} = 1.2546$ |
| 3 | *LOL1* | $\beta_{L,3} = 0.0508, B_{L,3} = 0.5048, c_{LOL} = -0.1108$ |
| 4 | *AT1G79150* | $\alpha_{L,4,1} = 0.8174, \alpha_{L,4,2} = -0.7887, \beta_{L,4} = 2.3792$<br>$B_{L,4} = 23.4766, c_{L,4} = 0.9019$ |
| 5 | *ANAC055* | $\alpha_{L,5,1} = 26.8670, \beta_{L,5} = 29.4074, B_{L,5} = 0.0493$<br>$c_{L,5} = 74.3481$ |
| 6 | *a-ERF-1* | $\beta_{L,6} = 0.2106, B_{L,6} = 1.9492, c_{L,6} = 0.8220$ |
| 7 | *ATML1* | $\alpha_{L,7,1} = -0.6137, \beta_{L,7} = 1.2522, B_{L,7} = 18.4637$<br>$c_{L,7} = 0.0043, \gamma_{L,7} = 0.5599$ |
| 8 | *CHE* | $\alpha_{L,8,1} = 19.9698, \alpha_{L,8,2} = 3.4368, \alpha_{L,8,3} = 20.0840$<br>$\beta_{L,8} = 39.8432, B_{L,8} = 13.822, \gamma_{L,8} = 18.3422$ |
| 9 | *RAP2.6L* | $\alpha_{L,9,1} = 0.4282, \beta_{L,9} = 0.7130, B_{L,9} = 2.8232$<br>$c_{L,9} = 0.0412$ |

**TABLE S2:** Estimated parameters of 9GRN using standard S-System model

| Number | Gene | Parameter |
| --- | --- | --- |
| 1 | *ORA59* | $\alpha_{SS,1} = 1.4005, g_{SS,1,1} = 0.8899, g_{SS,1,2} = -0.0087$<br>$g_{SS,1,3} = 0.3271, \beta_{SS,1} = 2.1743$ |
| 2 | *MYB51* | $\alpha_{SS,2} = 0.5605, g_{SS,2,1} = 0.0264, \beta_{SS,2} = 0.0531$ |
| 3 | *LOL1* | $\alpha_{SS,3} = 1.6419, g_{SS,3,1} = -0.0294, \beta_{SS,3} = 0.1880$ |
| 4 | *AT1G79150* | $\alpha_{SS,4} = 26.3739, g_{SS,4,1} = 0.3699, g_{SS,4,2} = -0.1299$<br>$g_{SS,4,3} = 0.0029, \beta_{SS,4} = 4.5064$ |
| 5 | *ANAC055* | $\alpha_{SS,5} = 0.7015, g_{SS,5,1} = 1.4538, g_{SS,5,2} = 0.0162$<br>$\beta_{SS,5} = 1.8916$ |
| 6 | *a-ERF-1* | $\alpha_{SS,6} = 0.5229, g_{SS,6,1} = 0.2218, \beta_{SS,6} = 0.0194$ |
| 7 | *ATML1* | $\alpha_{SS,7} = 26.6802, g_{SS,7,1} = -0.4408, g_{SS,7,2} = 0.0106$<br>$g_{SS,7,3} = 0.0063, \beta_{SS,7} = 0.9128$ |
| 8 | *CHE* | $\alpha_{SS,8} = 1.4648, g_{SS,8,1} = 0.1150, g_{SS,8,2} = 0.8517$<br>$g_{SS,8,3} = 0.1936, g_{SS,8,4} = 0.0145, \beta_{SS,8} = 1.9145$ |
| 9 | *RAP2.6L* | $\alpha_{SS,9} = 2.0164, g_{SS,9,1} = 0.5845, g_{SS,9,2} = 0.0058$<br>$\beta_{SS,9} = 0.7586$ |

**TABLE S3:** Estimated parameters of 9GRN using extended S-System model

| Number | Gene | Parameter |
| --- | --- | --- |
| 1 | *ORA59* | $\alpha_{ES,1} = 1.0174, g_{ES,1,1} = 0.6482, g_{ES,1,2} = -0.2417$<br>$g_{ES,1,3} = 0.8272, \beta_{ES,1} = 1.8866, \gamma_{ES,1,1} = 1.0577$ |
| 2 | *MYB51* | $\beta_{ES,2} = 1.0694, \gamma_{ES,2,1} = 9.0585, \gamma_{ES,2,2} = 1.9728$ |
| 3 | *LOL1* | $\beta_{ES,3} = 0.0498, \gamma_{ES,3,1} = 0.4994, \gamma_{ES,3,2} = -0.1328$ |
| 4 | *AT1G79150* | $\alpha_{ES,4} = 3.2664, g_{ES,4,1} = 0.7837, g_{ES,4,2} = 0.0692$<br>$\beta_{ES,4} = 2.3804, \gamma_{ES,4,1} = -0.7803, \gamma_{ES,4,2} = 0.1912$ |
| 5 | *ANAC055* | $\alpha_{ES,5} = 1.0481, g_{ES,5,1} = 1.0618, \beta_{ES,5} = 1.0110$<br>$\gamma_{ES,5,1} = -2.5859, \gamma_{ES,5,2} = 2.3646$ |
| 6 | *a-ERF-1* | $\beta_{ES,6} = 0.2740, \gamma_{ES,6,1} = 2.5325, \gamma_{ES,6,2} = 1.0798$ |
| 7 | *ATML1* | $\alpha_{ES,7} = 29.7754, g_{ES,7,1} = -0.3429, \beta_{ES,7} = 0.6780$<br>$\gamma_{ES,7,1} = -6.6253, \gamma_{ES,7,2} = 0.0047, \gamma_{ES,7,3} = 0.0074$ |
| 8 | *CHE* | $\alpha_{ES,8} = 1.7845, g_{ES,8,1} = 0.1761, g_{ES,8,2} = 0.0675$<br>$g_{ES,8,3} = 0.7054, \beta_{ES,8} = 1.6321, \gamma_{ES,8,1} = 2.1995$<br>$\gamma_{ES,8,2} = 0.7091$ |
| 9 | *RAP2.6L* | $\alpha_{ES,9} = 1.6086, g_{ES,9,1} = 0.6335, \beta_{ES,9} = 0.8880$<br>$\gamma_{ES,9} = 1.9096, \gamma_{ES,9,2} = 0.2511$ |

**TABLE S4:**
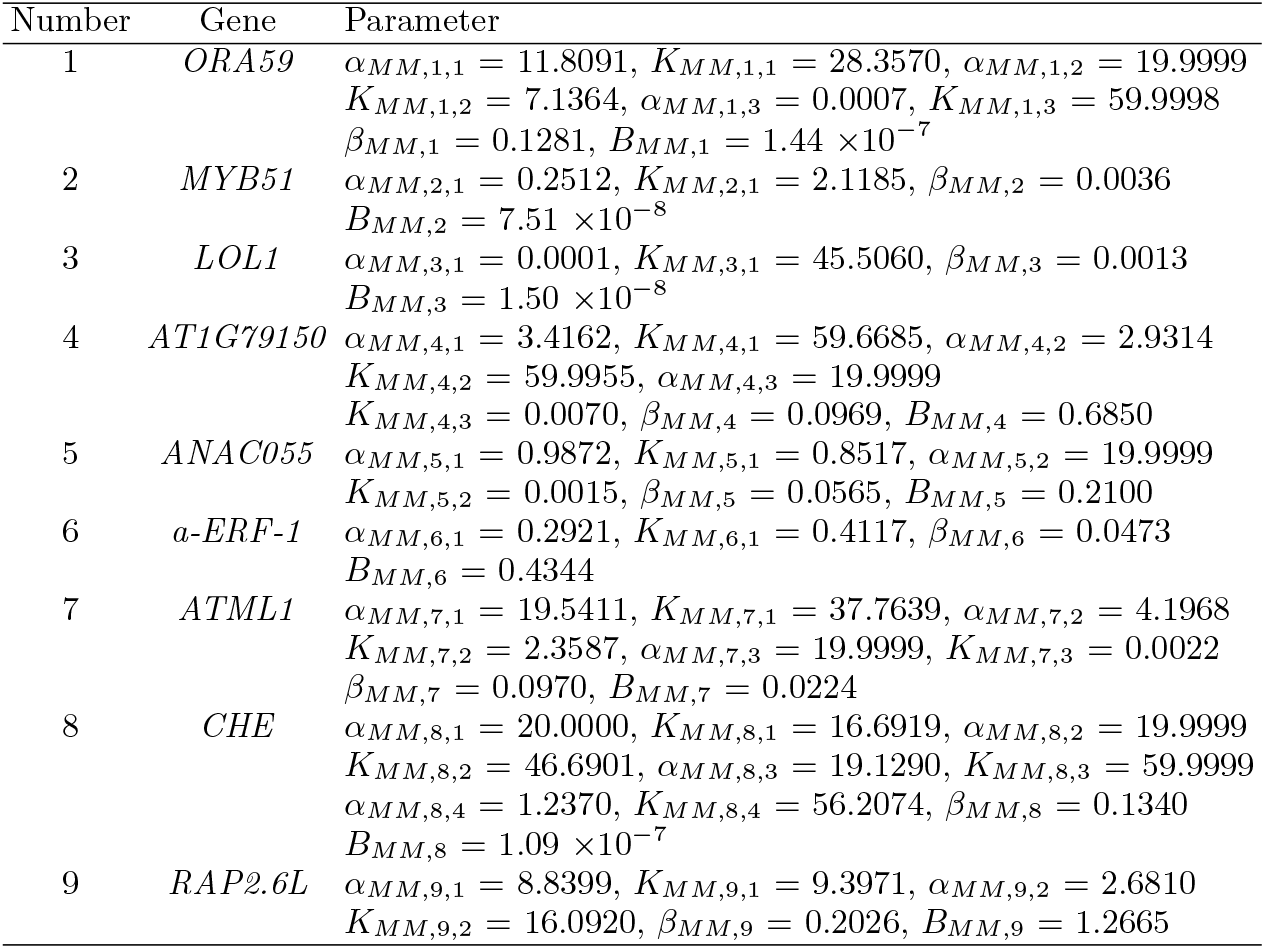
Estimated parameters of 9GRN using Michaelis-Menten model

| Number | Gene | Parameter |
| --- | --- | --- |
| 1 | *ORA59* | $\alpha_{MM,1,1} = 11.8091, K_{MM,1,1} = 28.3570, \alpha_{MM,1,2} = 19.9999$<br>$K_{MM,1,2} = 7.1364, \alpha_{MM,1,3} = 0.0007, K_{MM,1,3} = 59.9998$<br>$\beta_{MM,1} = 0.1281, B_{MM,1} = 1.44 \times 10^{-7}$ |
| 2 | *MYB51* | $\alpha_{MM,2,1} = 0.2512, K_{MM,2,1} = 2.1185, \beta_{MM,2} = 0.0036$<br>$B_{MM,2} = 7.51 \times 10^{-8}$ |
| 3 | *LOL1* | $\alpha_{MM,3,1} = 0.0001, K_{MM,3,1} = 45.5060, \beta_{MM,3} = 0.0013$<br>$B_{MM,3} = 1.50 \times 10^{-8}$ |
| 4 | *AT1G79150* | $\alpha_{MM,4,1} = 3.4162, K_{MM,4,1} = 59.6685, \alpha_{MM,4,2} = 2.9314$<br>$K_{MM,4,2} = 59.9955, \alpha_{MM,4,3} = 19.9999$<br>$K_{MM,4,3} = 0.0070, \beta_{MM,4} = 0.0969, B_{MM,4} = 0.6850$ |
| 5 | *ANAC055* | $\alpha_{MM,5,1} = 0.9872, K_{MM,5,1} = 0.8517, \alpha_{MM,5,2} = 19.9999$<br>$K_{MM,5,2} = 0.0015, \beta_{MM,5} = 0.0565, B_{MM,5} = 0.2100$ |
| 6 | *a-ERF-1* | $\alpha_{MM,6,1} = 0.2921, K_{MM,6,1} = 0.4117, \beta_{MM,6} = 0.0473$<br>$B_{MM,6} = 0.4344$ |
| 7 | *ATML1* | $\alpha_{MM,7,1} = 19.5411, K_{MM,7,1} = 37.7639, \alpha_{MM,7,2} = 4.1968$<br>$K_{MM,7,2} = 2.3587, \alpha_{MM,7,3} = 19.9999, K_{MM,7,3} = 0.0022$<br>$\beta_{MM,7} = 0.0970, B_{MM,7} = 0.0224$ |
| 8 | *CHE* | $\alpha_{MM,8,1} = 20.0000, K_{MM,8,1} = 16.6919, \alpha_{MM,8,2} = 19.9999$<br>$K_{MM,8,2} = 46.6901, \alpha_{MM,8,3} = 19.1290, K_{MM,8,3} = 59.9999$<br>$\alpha_{MM,8,4} = 1.2370, K_{MM,8,4} = 56.2074, \beta_{MM,8} = 0.1340$<br>$B_{MM,8} = 1.09 \times 10^{-7}$ |
| 9 | *RAP2.6L* | $\alpha_{MM,9,1} = 8.8399, K_{MM,9,1} = 9.3971, \alpha_{MM,9,2} = 2.6810$<br>$K_{MM,9,2} = 16.0920, \beta_{MM,9} = 0.2026, B_{MM,9} = 1.2665$ |

**TABLE S5:** WMSE for each individual in 9GRN using the data that is used in parameter estimation exercise. The sum of the individual WMSE yields the average total WMSE given in Table 3 in the main text.

| Model | Gene | WMSE | Gene | WMSE | Gene | WMSE |
| --- | --- | --- | --- | --- | --- | --- |
| Linear | *ORA59* | 0.00488 | *MYB51* | 0.00274 | *LOL1* | 0.00300 |
|  | *AT1G79150* | 0.00110 | *ANAC055* | 0.00441 | *a-ERF-1* | 0.00239 |
|  | *ATML1* | 0.00132 | *CHE* | 0.00228 | *RAP2.6L* | 0.00199 |
| Michaelis-Menten | *ORA59* | 0.01666 | *MYB51* | 0.00852 | *LOL1* | 0.00412 |
|  | *AT1G79150* | 0.00186 | *ANAC055* | 0.00697 | *a-ERF-1* | 0.00550 |
|  | *ATML1* | 0.00228 | *CHE* | 0.00433 | *RAP2.6L* | 0.00382 |
| Standard S-System | *ORA59* | 0.00548 | *MYB51* | 0.00396 | *LOL1* | 0.00354 |
|  | *AT1G79150* | 0.00165 | *ANAC055* | 0.00493 | *a-ERF-1* | 0.00428 |
|  | *ATML1* | 0.00258 | *CHE* | 0.00348 | *RAP2.6L* | 0.00244 |
| Extended S-System | *ORA59* | 0.00468 | *MYB51* | 0.00268 | *LOL1* | 0.00297 |
|  | *AT1G79150* | 0.00122 | *ANAC055* | 0.00363 | *a-ERF-1* | 0.00229 |
|  | *ATML1* | 0.00122 | *CHE* | 0.00249 | *RAP2.6L* | 0.00181 |

**TABLE S6:** WMSE for each individual in 9GRN using the data that is not used in parameter estimation exercise. The sum of the individual WMSE yields the average total WMSE given in Table 3 in the main text.

| Model | Gene | WMSE | Gene | WMSE | Gene | WMSE |
| --- | --- | --- | --- | --- | --- | --- |
| Linear | *ORA59* | 0.01145 | *MYB51* | 0.00499 | *LOL1* | 0.00466 |
|  | *AT1G79150* | 0.00223 | *ANAC055* | 0.00960 | *a-ERF-1* | 0.00554 |
|  | *ATML1* | 0.00298 | *CHE* | 0.00354 | *RAP2.6L* | 0.00385 |
| Michaelis-Menten | *ORA59* | 0.01526 | *MYB51* | 0.00739 | *LOL1* | 0.00676 |
|  | *AT1G79150* | 0.00263 | *ANAC055* | 0.01298 | *a-ERF-1* | 0.00804 |
|  | *ATML1* | 0.00387 | *CHE* | 0.00531 | *RAP2.6L* | 0.00688 |
| Standard S-System | *ORA59* | 0.01269 | *MYB51* | 0.00527 | *LOL1* | 0.00497 |
|  | *AT1G79150* | 0.00250 | *ANAC055* | 0.00987 | *a-ERF-1* | 0.00649 |
|  | *ATML1* | 0.00421 | *CHE* | 0.00441 | *RAP2.6L* | 0.00410 |
| Extended S-System | *ORA59* | 0.01130 | *MYB51* | 0.00477 | *LOL1* | 0.00456 |
|  | *AT1G79150* | 0.00253 | *ANAC055* | 0.00834 | *a-ERF-1* | 0.00522 |
|  | *ATML1* | 0.00295 | *CHE* | 0.00316 | *RAP2.6L* | 0.00362 |

**TABLE S7:** The TMSE of mutant analysis calculated using Eq. (S1).

| Model | TMSE |
| --- | --- |
| Linear | 1404.83 |
| Michaelis-Menten | 1753.05 |
| Standard S-System | 1263.18 |
| Extended S-System | 1285.56 |

**TABLE S8:** Average total WMSE for both ‘training’ and ‘validation’ data sets for 9GRN, with alternate Michaelis-Menten model, which the regulation types follow the linear model

| Model | Average Total WMSE<br>(training) | Average Total WMSE<br>(validation) |
| --- | --- | --- |
| Linear | 0.00267 | 0.00543 |
| Michaelis-Menten | 0.00614 | 0.00784 |
| Alternate Michaelis-Menten | 0.00285 | 0.00531 |
| Standard S-System | 0.00359 | 0.00606 |
| Extended S-System | 0.00256 | 0.00516 |

**TABLE S9:** Ranking model fits to experimental data based on AIC for 9GRN with alternate Michaelis-Menten model, which the regulation types follow the linear model. The notation *L, MM, AMM, SS, ES*, denote the linear, Michaelis-Menten, Alternate Michaelis-Menten, Standard S-System and Extended S-System models, respectively. Here *n_G_* = 9, *n_T_* = 48, *K*_Θ_ is the number of parameters in the model, 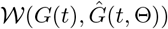 is the WMSE best fit to the data set used for parameter estimation, Δ_*i*_(AIC) is the AIC differences and *w_i_*(AIC) is the Akaike weights for each model.

| Model | *L* | *MM* | *AMM* | *SS* | *ES* |
| --- | --- | --- | --- | --- | --- |
| $K_\Theta$ | 38 | 58 | 58 | 38 | 44 |
| $\mathcal{W}(G(t), \hat{G}(t, \Theta))$ | 0.00267 | 0.00614 | 0.00285 | 0.00359 | 0.00256 |
| $\Delta_i(\text{AIC})$ | 8.182 | 397.506 | 74.266 | 134.696 | 0 |
| $w_i(\text{AIC})$ | 0.0164 | $4.14 \times 10^{-89}$ | $7.34 \times 10^{-17}$ | $5.55 \times 10^{-30}$ | 0.9836 |

**TABLE S10:** Estimated parameters of 9GRN using alternate Michaelis-Menten model, where the regulation types follow the linear model.

| Number | Gene | Parameter |
| --- | --- | --- |
| 1 | *ORA59* | $\alpha_{MM,1,1} = 41.5309, K_{MM,1,1} = 23.5156, \alpha_{MM,1,2} = 26.3578$<br>$K_{MM,1,2} = 6.1352, \alpha_{MM,1,3} = 39.5894, K_{MM,1,3} = 7.9278$<br>$\beta_{MM,1} = 3.1652, B_{MM,1} = 0.0595$ |
| 2 | *MYB51* | $\alpha_{MM,2,1} = 0.59623, K_{MM,2,1} = 0.3299, \beta_{MM,2} = 0.2967$<br>$B_{MM,2} = 2.5195$ |
| 3 | *LOL1* | $\alpha_{MM,3,1} = 1.7311, K_{MM,3,1} = 1.5207, \beta_{MM,3} = 0.1412$<br>$B_{MM,3} = 0.6472$ |
| 4 | *AT1G79150* | $\alpha_{MM,4,1} = 23.2089, K_{MM,4,1} = 9.0294, \alpha_{MM,4,2} = 0.0090$<br>$K_{MM,4,2} = 14.9596, \alpha_{MM,4,3} = 0.1659$<br>$K_{MM,4,3} = 2.0759, \beta_{MM,4} = 1.8107, B_{MM,4} = 4.8233$ |
| 5 | *ANAC055* | $\alpha_{MM,5,1} = 197.1760, K_{MM,5,1} = 30.9143, \alpha_{MM,5,2} = 1.4599$<br>$K_{MM,5,2} = 0.3304, \beta_{MM,5} = 1.9775, B_{MM,5} = 0.8868$ |
| 6 | *a-ERF-1* | $\alpha_{MM,6,1} = 1.1915, K_{MM,6,1} = 0.6445, \beta_{MM,6} = 0.2183$<br>$B_{MM,6} = 2.0356$ |
| 7 | *ATML1* | $\alpha_{MM,7,1} = 356.8399, K_{MM,7,1} = 0.1801, \alpha_{MM,7,2} = 0.1556$<br>$K_{MM,7,2} = 1.3908, \alpha_{MM,7,3} = 1.2922, K_{MM,7,3} = 1.9739$<br>$\beta_{MM,7} = 0.9991, B_{MM,7} = 6.2548$ |
| 8 | *CHE* | $\alpha_{MM,8,1} = 76.5103, K_{MM,8,1} = 27.8720, \alpha_{MM,8,2} = 72.6592$<br>$K_{MM,8,2} = 29.2936, \alpha_{MM,8,3} = 54.2288, K_{MM,8,3} = 30.6393$<br>$\alpha_{MM,8,4} = 0.4525, K_{MM,8,4} = 1.4687, \beta_{MM,8} = 2.6524$<br>$B_{MM,8} = 8.7780$ |
| 9 | *RAP2.6L* | $\alpha_{MM,9,1} = 17.8568, K_{MM,9,1} = 20.3954, \alpha_{MM,9,2} = 0.6174$<br>$K_{MM,9,2} = 0.8200, \beta_{MM,9} = 1.1531, B_{MM,9} = 7.8983$ |

## References

[1] B. Williamson, B. Tudzynski, P. Tudzynski, and J A L. Van Kan. Botrytis cinerea: the cause of grey mould disease. Molecular Plant Pathology, 8(5):561–580, 2007.

[2] Y. Jamir, M. Guo, H S. Oh, T. Petnicki-Ocwieja, S. Chen, X. Tang, M B. Dickman, A. Collmer, and J R. Alfano. Identification of Pseudomonas syringae type III effectors that can suppress programmed cell death in plants and yeast. Plant Journal, 37(4):554–565, 2007.

[3] A. Weiberg, M. Wang, F M. Lin, H. Zhao, I. Kaloshian, H D. Huang, and H. Jin. Fungal small RNAs suppress plant immunity by hijacking host RNA interference pathways. Science, 342(6154):118–123, 2013.

[4] J D G. Jones and J L Dangl. The plant immune system. Nature, 444(7117):323–329, 2006.

[5] SK Aoki, G Lillacci, A Gupta, A Baumschlager, D Schweingruber, and M Khammash. A universal biomolecular integral feedback controller for robust perfect adaptation. Nature, 570:533–537, 2019.

[6] M. Foo, I. Gherman, P. Zhang, D G. Bates, and K J. Denby. A framework for engineering stress resilient plants using genetic feedback control and regulatory network rewiring. ACS Synthetic Biology, 7:1553–1564, 2018.

[7] Danny W-K Ng, Jayami K Abeysinghe, and Maedeh Kamali. Regulating the regulators: the control of transcription factors in plant defense signaling. International Journal of Molecular Sciences, 19:3737, 2018.

[8] Monika Sood, Dhriti Kapoor, Vipul Kumar, Namarta Kalia, Renu Bhardwaj, Gagan PS Sidhu, and Anket Sharma. Mechanisms of plant defense under pathogen stress: a review. Current Protein and Peptide Science, 22:376–395, 2021.

[9] Iulia Gherman. Engineering stress resilient plants using gene regulatory network rewiring. PhD thesis, University of Warwick, 2018.

[10] O. Windram, P. Madhou, S. McHattie, C. Hill, R. Hickman, E. Cooke, D J. Jenkins, C A. Penfold, L. Baxter, E. Breeze, S J. Kiddle, J. Rhodes, S. Atwell, D J. Kliebenstein, Y S. Kim, O. Stegle, K. Borgwardt, C. Zhang, A. Tabrett, R. Legaie, J. Moore, B. Finkenstadt, D L. Wild, A. Mead, D. Rand, J. Beynon, S. Ott, V. Buchanan-Wollaston, and K J. Denby. *Arabdiopsis* defense against *Botrytis cinerea*: chronology and regulation deciphered by high-resolution temporal transcriptomic analysis. Plant Cell, 24:3530–3557, 2012.

[11] T Schlitt and A Brazma. Current approaches to gene regulatory network modelling. BMC Bioinformatics, 8:S9, 2009.

[12] J Tegner, MKS Yeung, J Hasty, and JJ Collins. Reverse engineering gene networks: integrating genetic perturbations with dynamical modeling. Proceedings of the National Academy of Sciences, 100:5944–5949, 2003.

[13] H Hache, H Lehrach, and R Herwiga. Reverse engineering of gene regulatory networks: a comparative study. EURASIP Journal on Bioinformatics and Systems Biology, 2009:1–12, 2009.

[14] Christopher A. Penfold and David L. Wild. How to infer gene networks from expression profiles, revisited. Interface Focus, 1(6):857–870, 2011.

[15] Lian En Chai, Swee Kuan Loh, Swee Thing Low, Mohd Saberi Mohamad, Safaai Deris, and Zalmiyah Zakaria. A review on the computational approaches for gene regulatory network construction. Computers in Biology and Medicine, 48:55–65, 2014.

[16] Alejandro F Villaverde, John Ross, and Julio R Banga. Reverse engineering cellular networks with information theoretic methods. Cells, 2(2):306–329, 2013.

[17] Veronica Vinciotti, Luigi Augugliaro, Antonino Abbruzzo, and Ernst C Wit. Model selection for factorial gaussian graphical models with an application to dynamic regulatory networks. Statistical Applications in Genetics and Molecular Biology, 15(3):193–212, 2016.

[18] F Emmert-Streib, GV Glazko, G Altay, and R de Matos Simoes. Statistical inference and reverse engineering of gene regulatory networks from observational expression data. Frontiers in Genetics, 3:8, 2012.

[19] Fei He, Ettore Murabito, and Hans V Westerhoff. Synthetic biology and regulatory networks: where metabolic systems biology meets control engineering. Journal of the Royal Society Interface, 13(117):20151046, 2016.

[20] TA Long, SM Brady, and PN Benfey. Systems approaches to identifying gene regulatory networks in plants. Annual Review of Cell and Developmental Biology, 24:81–103, 2008.

[21] LV Den Broeck, M Gordon, D Inze, C Williams, and R Sozzani. Gene regulatory network inference: connecting plant biology and mathematical modeling. Frontiers in Genetics, 11:457, 2020.

[22] C.C.N. Wang, P.-C. Chang, K.-L. Ng, C.-M. Chang, P.C.Y. Sheu, and J.J.P Tsai. A model comparison study of the flowering time regulatory network in *Arabidopsis*. BMC Systems Biology, 8(15), 2014.

[23] G Karlebach and R Shamir. Modelling and analysis of gene regulatory networks. Nature Reviews Molecular Cell Biology, 9:770–780, 2008.

[24] AR Chowdhury and M Chetty. Reconstruction of large-scale gene regulatory network using S-System model. Evolutionary Computation in Gene Regulatory Network Research, 2016:185–210, 2016.

[25] M. Foo, D G. Bates, and O E. Akman. A simplified modelling framework facilitates more complex representations of plant circadian clocks. PLoS Computational Biology, 16:e1007671, 2020.

[26] M A. Savageau. Biochemical systems analysis ii. the steady state solutions for an n-pool system using a power-law approximation. Journal of Theoretical Biology, 25:370–379, 1969.

[27] M A. Savageau. Design principles for elementary gene circuits: elements, methods, and examples. Chaos, 11:142–159, 2001.

[28] E O. Voit, H A. Martens, and S W. Omholt. 150 years of mass action law. PLoS Computational Biology, 11:e1004012, 2015.

[29] Y Maki, D Tominaga, M Okamoto, S Watanabe, and Y Eguchi. Development of a system for the inference of large scale genetic networks. Biocomputing, 2001:446–458, 2000.

[30] H Bolouri and EH Davidson. Modeling transcriptional regulatory networks. Bioessays, 24:1118–1129, 2002.

[31] P Rué and J Garcia-Ojalvo. Modeling gene expression in time and space. Annual Review of Biophysics, 42:605–627, 2013.

[32] ASK Youseph, M Chetty, and G Karmakar. Gene regulatory network inference using Michaelis-Menten kinetics. Proceedings of IEEE Congress of Evolutionary Computation, 25-28 May 2015, Sendai, Japan, pages 2392–2397, 2015.

[33] H Wang, L Qian, and E Dougherty. Inference of gene regulatory networks using S-system: a unified approach. IET Systems Biology, 4:145–156, 2010.

[34] Leander Dony, Fei He, and Michael P H Stumpf. Parametric and non-parametric gradient matching for network inference: a comparison. BMC Bioinformatics, 20(1):1–12, 2019.

[35] Ann C Babtie, Paul Kirk, and Michael P H Stumpf. Topological sensitivity analysis for systems biology. Proceedings of the National Academy of Sciences, 111(52):18507–18512, 2014.

[36] Tarmo Äijö and R Bonneau. Biophysically motivated regulatory network inference: progress and prospects. Human Heredity, 81:62–77, 2016.

[37] Michael M Saint-Antoine and Abhyudai Singh. Network inference in systems biology: recent developments, challenges, and applications. Current Opinion in Biotechnology, 63:89–98, 2020.

[38] K.P. Burnham and D.R. Anderson. Information and Likelihood Theory: A Practical Information-Theoretic Approach. Springer-Verlag, New York, 2002.

[39] K.P. Burnham and D.R. Anderson. Multimodel inference: Understanding AIC and BIC in model selection. Sociology Methods and Research, 33(2):261–304, 2004.

[40] H T. Banks and M L. Joyner. AIC under the framework of least squares estimation. Applied Mathematics Letters, 74:33–45, 2017.

[41] E.-J. Wagenmakers and S. Farrell. AIC model selection using akaike weights. Psychonomin Bulletin and Review, 11(1):192–196, 2004.

[42] GK Acker, AD Johnson, and MA Shea. Quantitative model for gene regulation by lambda phage repressor. Proccedings of the National Academy of Sciences, 79:1129–1133, 1982.

[43] MK Transtrum and P Qiu. Optimal experiment selection for parameter estimation in biological differential equation models. BMC Bioinformatics, 13:181, 2012.

[44] NMG Paulino, M Foo, J Kim, and DG Bates. Robustness analysis of a nucleic acid controller for a dynamic biomolecular process using the structured singular value. Journal of Process Control, 78:34–44, 2019.

[45] S Marino, IB Hogue, CJ Ray, and DE Kirschner. A methodology for performing global uncertainty and sensitivity analysis in systems biology. Journal of Theoretical Biology, 254:178–196, 2008.

[46] R Sheikholeslami and S Razavi. Progressive latin hypercube sampling: an efficient approach for robust sampling-based analysis of environmental models. Environmental Modelling and Software, 93:109–126, 2017.

[47] JBE Rinõn, RG Mendoza, and VMP Mendoza. Parameter estimation of an S-system model using hybrid genetic algorithm with the aid of sensitivity analysis. Proceedings of Philippines Computing Science Congress, Manila, Philippines, 28-30 March 2019, pages 94–102, 2019.

[48] JK Kim and JJ Tyson. Misuse of the Michaelis-Menten rate law for protein interaction networks and its remedy. PLoS Computational Biology, 16:e1008258, 2020.

[49] M. Foo, J. Kim, and D G. Bates. Modelling and control of gene regulatory networks for perturbation mitigation. IEEE/ACM Transactions on Computational Biology and Bioinformatics, 16:583–595, 2018.

